# Vicarious Somatotopic Maps Tile Visual Cortex

**DOI:** 10.1101/2024.10.21.619382

**Authors:** Nicholas Hedger, Thomas Naselaris, Kendrick Kay, Tomas Knapen

## Abstract

Our sensory systems work together to generate a cohesive experience of the world around us. Watching others being touched activates brain areas representing our own sense of touch: the visual system recruits touch-related computations to simulate bodily consequences of visual inputs. One long-standing question is how the brain implements this interface between visual and somatosensory representations. To address this question, we developed a method to simultaneously map somatosensory body part tuning and visual field tuning throughout the brain. Applying this method on ongoing co-activations during rest resulted in detailed maps of the body-part tuning in the brain’s endogenous somatotopic network. During movie watching, somatotopic tuning explains responses throughout the entire dorsolateral visual system, revealing an array of somatotopic body maps that tile the cortical surface. The tuning of these maps aligned with those of visual maps, and predicted both preferences for visual field locations and the visual-category preferences for body parts. These results reveal a mode of brain organization in which aligned visual-somatosensory topographic maps connect visual and bodily reference frames. This cross-modal interface is ideally situated to translate raw sensory impressions into more abstract formats useful for action, social cognition, and semantic processing.

## Main

In humans, input from one sensory modality can influence processing in other modalities to enrich perception and understanding. One striking example is how visual inputs recruit somatosensation, our sense of touch: when seeing others in pain, we may flinch, wince, and even remark that we ‘felt their pain’. Indeed, when observing others, our brain often responds ‘as if’ their tactile experience were our own: Human neuroimaging work has demonstrated extensive, body-part specific activations in motor and somatosensory cortex when participants view others acting on objects^1^ or being touched.^2^ This cross-modal relationship between vision and touch is bi-directional: regions of high-level visual cortex similarly exhibit body-part specific activations when participants perform unseen actions.^3, 4^ These “non-afferent”, or ‘foreign source” activations of our sensorimotor and visual systems have been linked to vicarious and empathic experiences.^5, 6^ Despite the pervasive nature of these inter-modal, non-afferent activations, we lack understanding of how the brain connects the computational machinery of vision and somatosensation.

Beyond non-afferent recruitment by visual inputs, our somatosensory system likely has modes of operation that are independent of any sensory input. Much of our spontaneous thought is self-referential, likely engaging unimodal somatosensory regions through endogenous, body-referenced processes like emotional introspection and interoception.^7, 8^ By contrast, when somatosensation is non-afferently recruited by exogenous visual inputs, our brain must recode visual signals into a simulated somatosensory representation. This implies involvement of multimodal neural sites with selective tuning tethered to both visual and somatosensory reference frames. This putative joint tuning to visual and bodily coordinates aligns with models of ‘embodied’ visual perception,^9^ which claim that visual perception is inherently multisensory and shaped by potential bodily interactions with observed stimuli. One prediction flowing from this idea is that neural responses to visual stimulation will be better explained by models incorporating selective tuning in both visual and somatosensory modalities, rather than visual tuning alone. Testing this set of normative predictions requires an explicit computational model that can capture multimodal tuning during both endogenous and non-afferent stimulation.

Somatosensory tuning in multimodal sites may relate to visual tuning in diverse ways, instantiating different multisensory computations. One hypothesis is that the organization of the brain’s visual and somatosensory tuning align to reflect environmental statistics. This may be implemented by imposing the visual system’s retinal reference frame on somatosensory tuning to align with regularities in ecological body part positions.^10^ For instance, for a given brain region, somatosensory tuning preferences for foot sensations may predict visual tuning preferences for the lower visual field locations wherein they typically appear. Another hypothesis is that tuning alignment between somatosensation and vision may also play out at a more category-selective level of visual function, such that, for instance, somatosensory selectivity for foot sensations predicts visual selectivity for presentations of feet, independent of their spatial location.

Here we show that modeling the topographic structure of brain activations uncovers the computational motifs connecting visual and touch representations and reveals distinct roles of somatosensation in endogenous thought and naturalistic vision. We show that responses in large swaths of cortex are predictable from structured connectivity to primary somatosensory cortex during rest. This connectivity extended into dorsolateral visual cortex during movie-watching, revealing multiple visually-evoked somatotopic maps. The tuning of these maps aligned with visual tuning along multiple dimensions: they predict preferences for visual field locations more dorsally and visual body-part preferences more ventrally. These visual-somatic topographic alignments across dorsolateral visual cortex may provide a common language for sharing structured information between sensory modalities, supporting cohesive perception and cognition.

### Endogenous connectivity with S1 reveals intrinsic somatotopic organization in parietal, frontal and insular cortex

We developed a computational model of joint visual and somatosensory tuning that capitalizes on a shared principle of our sensory systems: topographic organization. Primary visual cortex (V1) and primary somatosensory cortex (S1) contain retinotopic and somatotopic maps, with neighboring regions tuned to neighboring locations in the visual field^11^ or on the body,^12^ respectively (**Figure 1 a-b**). For every *target* voxel in the brain, we estimate the spatial patterns (connective fields) on the *source regions* of V1 and S1 that best explain its BOLD time-course (for details, see **methods**). Since patterns in these source regions relate directly to visual field and body positions, they allow connectivity-derived mapping of voxels’ visual and body-part tuning. In effect, the method leverages the profile of source-to-target connectivity to ‘project’ the topography of the source regions into target regions, revealing neurally-referred retinotopic and somatotopic maps (**Figure 1 c,d**).

**Figure 1.**
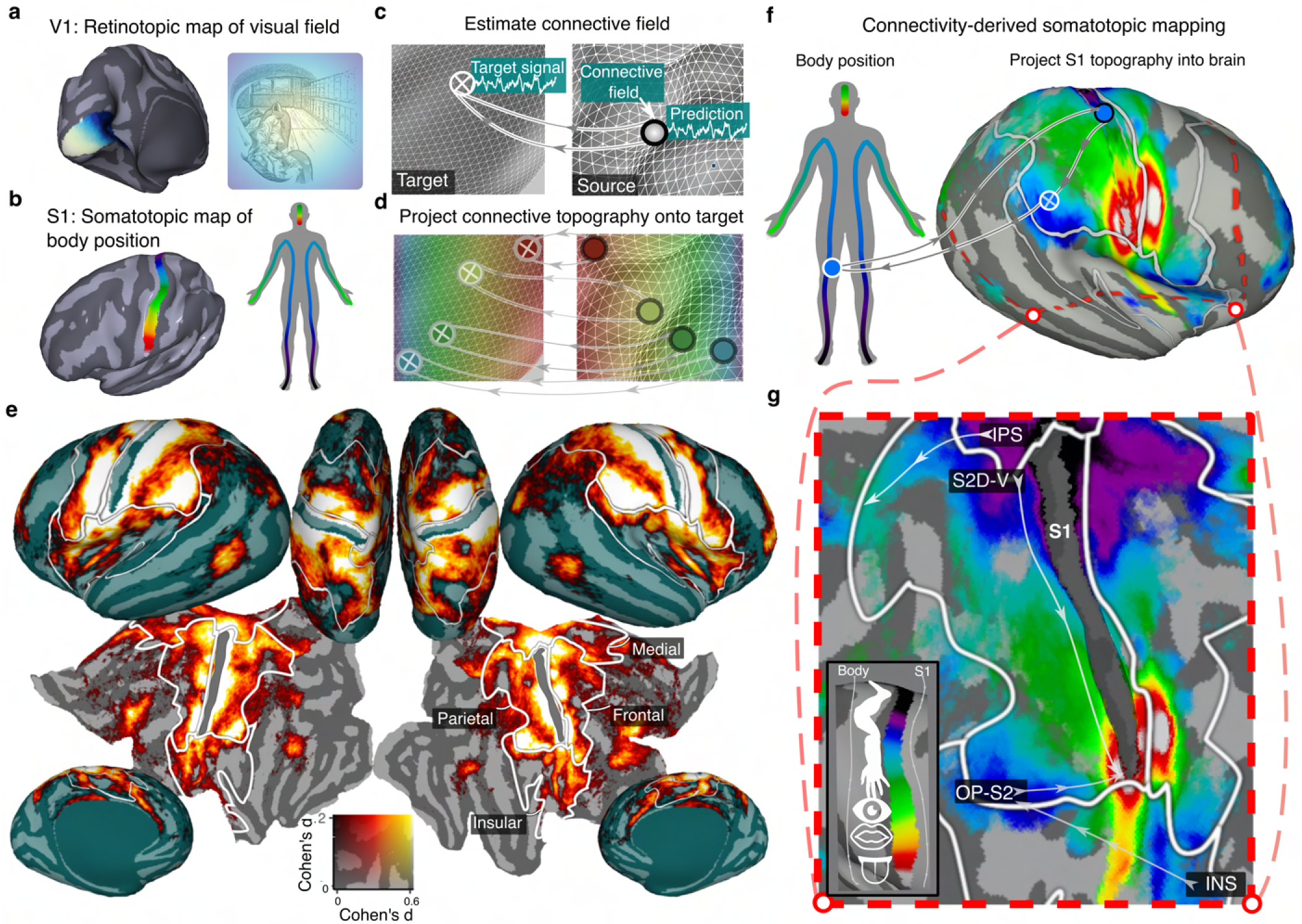
Connective field modeling reveals endogenous somatotopic network. **a-b.** Schematic representation of the retinotopic organization of human primary visual cortex (V1) and somatotopic organization of primary somatosensory cortex (S1). Neighboring positions in V1 and S1 are associated with sensitivity to neighboring positions in the visual field and body respectively (colors). **c.** Connective field modeling methodology. A time-varying response within a target region (e.g. higher level somatosensory cortex) is modeled as some combination of signals (i.e. a ‘connective field’) deriving from a source region (e.g. S1). **d.** With connective fields estimated for every part of the target region, the topographic map of the source can be projected onto the target according to the connectivity profile (colors). **e.** Depicts the strength of somatotopic connectivity during resting state on the cortical surface, expressed as Cohen’s d. White borders indicate the boundaries of gross-anatomical somatosensory regions denoted by the labels. **f.** Somatotopic mapping. Since positions in S1 correspond to sensations in positions on the body (colors), spatial connectivity with S1 can be leveraged to project preferred body position into the rest of the brain, revealing connectivity-derived somatotopic structure from resting-state data. **g.** Shows the same data for the cutout region in **f** demarcated by the red outline. Data are thresholded to show vertices with significant somatotopic connectivity. Arrows indicate the location of homuncular gradients described in the main text. The S1 source region (the data underlying the design matrix for our modeling) is removed so only target data are shown.

Recent work has revealed that the retinotopic organization of the visual system can be revealed from BOLD responses obtained in the absence of external stimulation (during rest).^13^ This suggests the brain has endogenous sensory-topographic structure that is recruited during endogenous thought. Here, we first tested whether our model can similarly reveal endogenous somatotopic structure from rest, which is typically revealed via exogenous tactile stimulation.^12^ To this end, we fit our model to one hour of 7T resting state data from each of 174 subjects from the Human Connectome Project (HCP).

**Figure 1 e** depicts the performance of S1 connective fields in predicting brain responses during rest, expressed as the degree of ‘somatotopic connectivity’ (**see Methods)**. A large swath of cortex, encompassing classically defined somatosensory regions in Parietal, Frontal, Medial and Insular cortex (**see Methods**)^12^ exhibited somatotopic connectivity (all *p<*10^−8^, min Cohen’s *d* = 0.41: **Figure 1 f-g**), with strongest effects detected in parietal and frontal cortex (Main effect of ROI: *F* (2.5,431.69) = 488.66, *p<*10^−125^).

With robust intrinsic somatotopic connectivity established, we next examined the underlying topographic structure of these activations by translating the connective field profiles into preferred body positions (**Figure 1f**), projecting the S1 somatotopic map into the brain. This reveals several orderly somatotopic gradients that mirror classical hallmarks of human somatosensory organization^14, 15^ (**Figure 1g**): one dorsal to ventral toes-tongue gradient along the central sulcus (*S2D-V*), one posterior-anterior from the parietal operculum/S2 (*OP-S2*), and a further double-gradient in insular cortex (*INS*). Lastly, another anterior-posterior gradient along the intraparietal sulcus (*IPS*) can also be observed. The somatotopic structure in each of the classically defined somatosensory regions was also replicated in independent subject splits (Parietal: M*z* =0.99, 95% CI = [0.97,1.02], Frontal: M*z* =1.08, 95% CI = [1.06,1.11], Medial: M*z* =0.59, 95% CI = [0.56,0.62], Insular: M*z* =0.81, 95% CI = [0.79,0.84]) (**Extended Data Figure E4)**.

A further known hallmark of somatotopic maps is their biased representations: over- and underrepresentation of certain body parts, putatively reflecting their respective functions.^12^ These hallmarks are also observed from our connectivity measures: medial regions had the largest proportion of lower-limb/trunk tuned locations (max *p* for pairwise comparisons =10**^-^**^28^), consistent with reports of stroke-induced lesions to this area in patients with leg-related pathology.^16^ By contrast, the largest proportion of upper-limb tuned locations were observed in parietal cortex (max *p* =.029), consistent with their proposed functional specialization for reaching and grasping actions.^17^

The anatomical loci, extents and detailed tuning properties of the somatotopic maps revealed by our model mirror those obtained from studies that employ tactile or motor interactions.^12, 15^ Crucially, however, our model demonstrates the intrinsic nature of this organization by establishing these principles under conditions devoid of external sensorimotor stimulation.

### Movie-watching extends somatotopic network into classically ‘visual’ cortex

Our analysis of resting state activity revealed that intrinsic somatotopic organization is present across a large extent of cortex and shares the structure expected from activity evoked by somatosensory stimulation. To ask how our somatotopic network is influenced by non-afferent naturalistic visual inputs, we next performed our connective field modeling on one hour of 7T data obtained from the same participants during movie watching.

**Figure 2a** depicts the difference in somatotopic connectivity strength between rest and movie-watching. In regions abutting S1, such as primary motor cortex (M1) and Broadman area 2, we see similar levels of somatotopic connectivity during movie-watching and resting state (*t* (173) =.96, *p*=.966, *t* (173) =-.54, *p*=.591). Moreover, the somatotopic organization of the network identified during rest was similar during movie watching **(Extended Data Figure E5)**. However, relative to rest, a much larger swathe of cortex (50%, 95% CI = [46%, 55%]) encompassing regions beyond our endogenous resting-state network exhibits somatotopic connectivity during movie-watching (**Figure 2b,c**).

**Figure 2.**
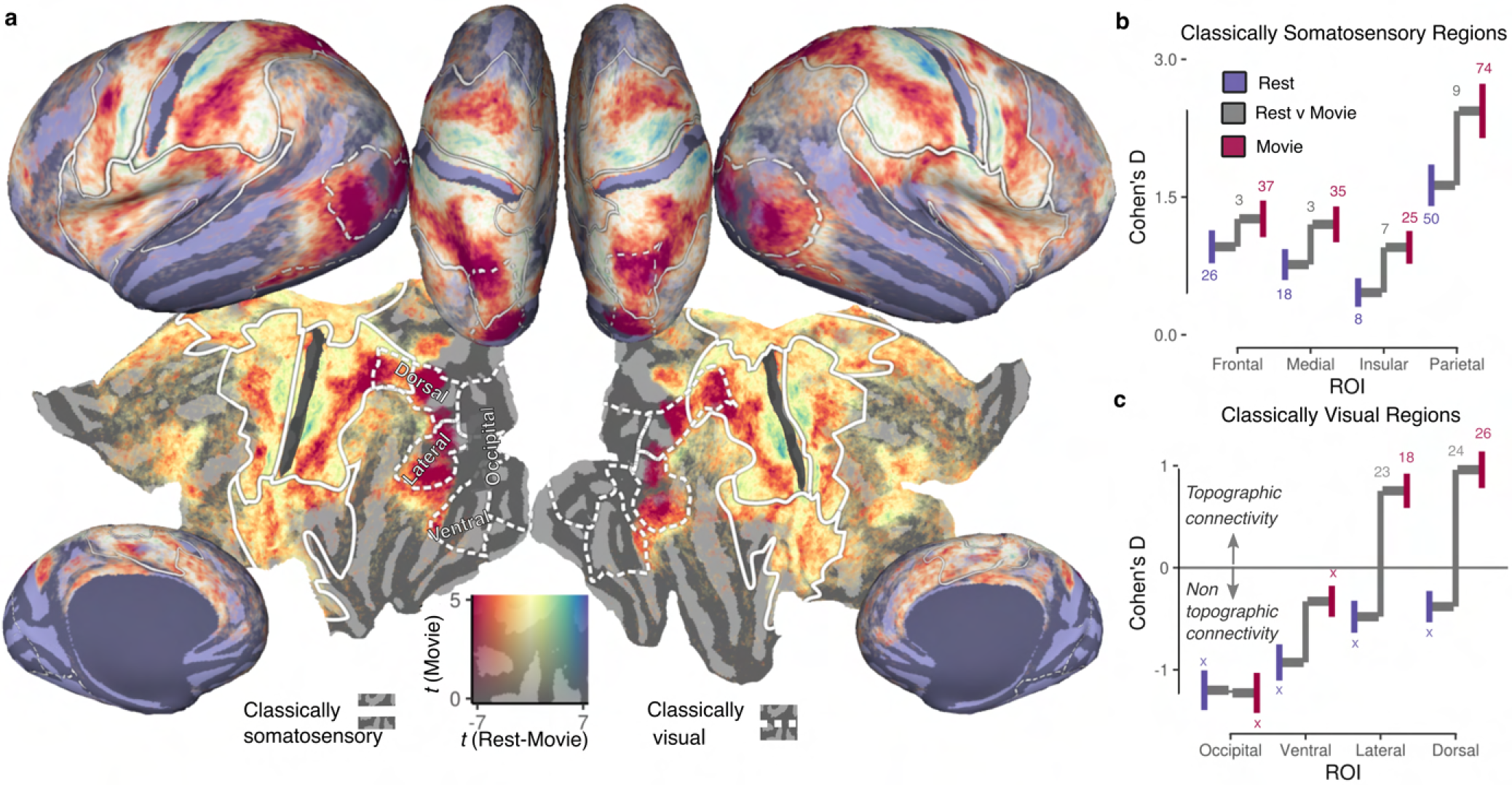
Somatotopic connectivity in visual cortex during movie watching. **a.** Contrasts somatotopic connectivity between rest and movie (red= greater during movie, blue=greater during rest). Solid overlays indicate the somatotopic ROIS outlined previously, dotted lines indicate gross divisions of conventionally-defined visual cortex (see **Methods**). **b.** Throughout the somatosensory network, movie watching increases the strength of somatosensotopic connectivity. Colored Vertical lines indicate the 95% confidence range of the estimate. Connecting grey lines illustrate the difference between conditions. Grey and colored numbers indicate the −log10 transformed p value for the one-sample difference from 0 and the paired difference between task conditions respectively. **c.** Focusing on the increased somatotopic connectivity in the visual system, we find that this connectivity is increased selectively in the lateral and dorsal portions of the visual system.

Critically, the additional somatotopic territory revealed by movie-watching encompassed even classically-defined ‘visual’ regions (**see Methods)**, which show increased somatotopic connectivity relative to rest (Main effect of task: *F* (1,173)=132.26, *p<*10**^-^**^22^) (**Figure 2c**). Dorsal and lateral visual regions specifically showed strongest increases (*p<*10**^-^**^23^), consistent with the proposed roles of these processing streams in ‘vision for action’ and the dynamic aspects of social perception,^18^ respectively (Interaction between stream and task: *F* (1.71, 296.48)=27.73, *p<*10**^-^**^9^).

### Dorsolateral visual responses balance retinotopic and somatotopic activations

Our analyses of movie-waching data reveal somatotopically structured activations in extrastriate cortex. This raises the natural question of how they relate to and coincide with the retinotopic activations more traditionally associated with these cortical areas. Since our analysis simultaneously estimates somatosensory and visual connective fields and performs automatic variance partitioning (**see Extended Data Figure E1a-f**), we can quantify the relative importance of somatotopic and retinotopic activations in explaining BOLD responses, thereby explicitly testing the centrality of somatotopic responses in the visual system.

**Figure 3a-f** depicts the balance of explained variance by visual (blue) and somatosensory (red) connective fields in the average-participant time series. Occipital visual cortex colors strongly blue and the endogenous somatotopic network centered on the central sulcus colors strongly red - indicating strongly unimodal activation patterns. Placed between these purely retinotopic or somatotopic regions, our analysis reveals a band of dorsolateral visual cortex with systematic, bilaterally symmetric alternations in the strength of somatotopic and retinotopic connectivity (see rectangular cutout/inset in **Figure 3b**.

**Figure 3.**
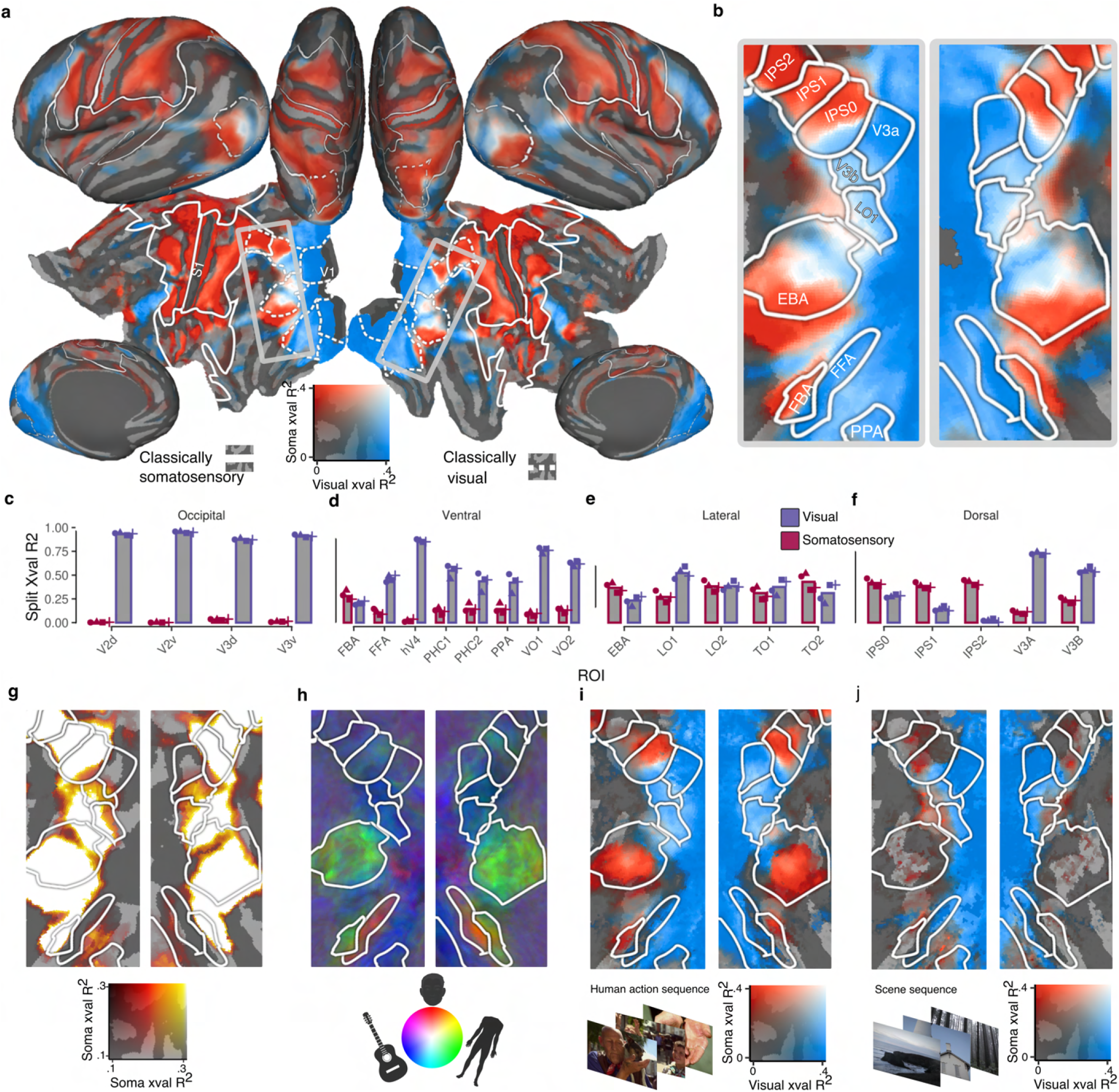
Multimodal topographic connectivity in dorsolateral visual cortex. **a.** Connective field model performance from each source region (V1 - blue, S1 - red) quantified by cross-validated explained variance. Solid and dashed overlays indicate the somatotopic ROIS and conventionally-defined visual regions defined in ( Figure 2). The rectangle marks a band of extrastriate cortex depicted in ensuing panels. **b.** Zoomed portion of extrastriate cortex in which preferred connective field modality reveals bilaterally symmetric alternations. **c-f.** Bar charts depict region-based average of S1 and V1-connective field derived cross-validated variance explained. In this average-participant representation, several regions in traditionally defined visual cortex (FBA, EBA, TO2, and IPS0-2) show a somatosensory-dominant connectivity profile. Symbols indicate estimates from movie and subject half-splits of the data. **g.** Depicts unimodal (somatotopic) cross validated variance explained. In the dorsal and lateral streams, V3A was the only region with no somatotopic connectivity. Variance explained drops off ventrally to FBA and posteriorly to LO1/V3B where retinotopic responses dominate. **h.** The RGB colormap depicts visual selectivity for faces, bodies and objects (peak-normalized t statistics) from an independent functional localiser. EBA and FBA appear as two green regions in lateral and ventral visual cortex, overlapping areas with dominant somatotopic connectivity in **b**. **i.** The same outcomes as in **b** are shown, estimated from a movie sequence involving human <u>actions and in **j.** estimated from a movie sequence depicting a movement through scenes, with no humans present.</u>

Although far-flung from classical somatosensory regions, much of this region shown in **Figure 3b** exhibits multimodal topographic connectivity (i.e. both retinotopic and somatotopic) during movie-watching. Taking the extrastriate body area (EBA) as an example, our model reveals clear alternations in modality preference that may provide a clarifying and unifying way of understanding the diversity of functions attributed to this large region.^19^ Specifically, the more posterior-dorsal portion, which we find to be primarily retinotopically-tuned (blue), overlaps with atlas definitions of the retinotopic regions TO and LO2 and terminates at their anterior boundary^20^, while the more anterior-ventral, somatotopically-tuned (red) portion overlaps with atlas definitions of motion-sensitive MT^21^ (see **see Extended Data Figure E7**) and extends anteriorly beyond EBA, consistent with the claimed existence of an ‘action-related region’ partially overlapping with EBA.^22^ Moreover, EBA is one of several dorsolateral ‘visual’ regions whose signal variance is best explained by topographic connectivity with S1 rather than V1 (**Figure 3b-f**).

Across subjects, all lateral and dorsal ROIs of the visual system (with the exception of V3A) exhibited multimodal topographic connectivity (max somatotopic *p* =.024, max retinotopic *p <*10^−22^) whereas in ventral regions, this multimodal connectivity was unique to the fusiform body area (FBA) (retinotopic *p<*10^−22^, somatotopic *p<*10^−8^) - a distinctive functional profile among ventral visual regions that sharply delineates FBA from neighboring FFA, as well as ventral retinotopic regions (VO, PHC) which only exhibit retinotopic activations. These findings establish that somatotopic processing may be unexpectedly pivotal in the visual system’s responses to dynamic naturalistic stimulation in the lateral and dorsal, but not the ventral visual system.

### Vicarious Somatotopic responses in both EBA and FBA

What aspects of movie-watching drive these somatotopic responses in dorsolateral visual cortex? Whilst movie-watching introduces exogenous stimulation relative to resting-state, the eliciting stimulus for the observed somatotopic responses is nonetheless *visual* - there is no afferent sensorimotor stimulation. Indeed, further analyses with M1 as an additional, third source region found that S1 connective fields greatly outperformed M1 in explaining dorsolateral responses during movie watching (**Extended Data Figure E6**). This superiority points to the primacy of non-afferent somatosensation over simulated or executed motor programs in explaining signal variance^23^

A striking feature of the data in **Figure 3a-b** is that the location of predominant somatotopic responses overlap substantially with known locations of two regions with robust preferences for *visual* presentations of bodies: the Extrastriate Body Area and Fusiform Body Area (FBA).^24^ Indeed, referencing our outcomes against independent functional localizer data (**Figure 3h**) indicated that somatotopic responses in visual cortex were more closely associated with visual selectivity for bodies than selectivity for faces (*p<*10^−4^), objects (*p* =.012) or places (*p* =.028), pointing to a visual-category-selective somatotopic response in these regions. To verify that indeed human observation drives somatotopic responses in visual cortex, we repeated our analysis on two sections of the movie: one that depicts human agents interacting with objects and one where the camera pans through various environmental locations with no human agents present. This revealed that the somatotopic responses in EBA and FBA were indeed contingent on viewing human agents, with both regions exhibiting greater somatotopic connectivity during the human sequence (*max p<*10^−2^) (**Figure 3i-j**). This supports the notion that these somatotopic responses reflect vicarious and selective responses to the human actions depicted in the movie, rather than generic responses to visual input, or bodily engagement with camera movements.^25^

### Dorsolateral visual cortex is tiled with somatotopic maps

Somatotopic connectivity explains a large proportion of BOLD signal variance across dorsolateral visual cortex during movie watching. This raises the intriguing possibility that, as in the core somatosensory network across frontal and parietal lobes, visual cortex’ body-part tuning may be structured as orderly, somatotopic map-like arrangements on the cortical surface. To test this possibility, we projected connective-field estimated somatotopic tuning onto this region. This revealed multiple gradients of connectivity-derived body-part tuning separated by reversals, a canonical signature of the presence of cortical field maps (**Figure 4a**). We tentatively outlined these maps as quadrilaterals on the cortical surface, and used projective transformation to render them into rectangles, concatenating them along the principal axis of the gradient. This renders each hemisphere’s somatotopic maps into a common space. This somatotopic structure, shown in **Figure 4b**, was highly consistent between hemispheres both across maps (*r* =.88, p*<*10^−16^, **Figure 4c**) and on an individual map level basis (**Figure 4e**).

**Figure 4.**
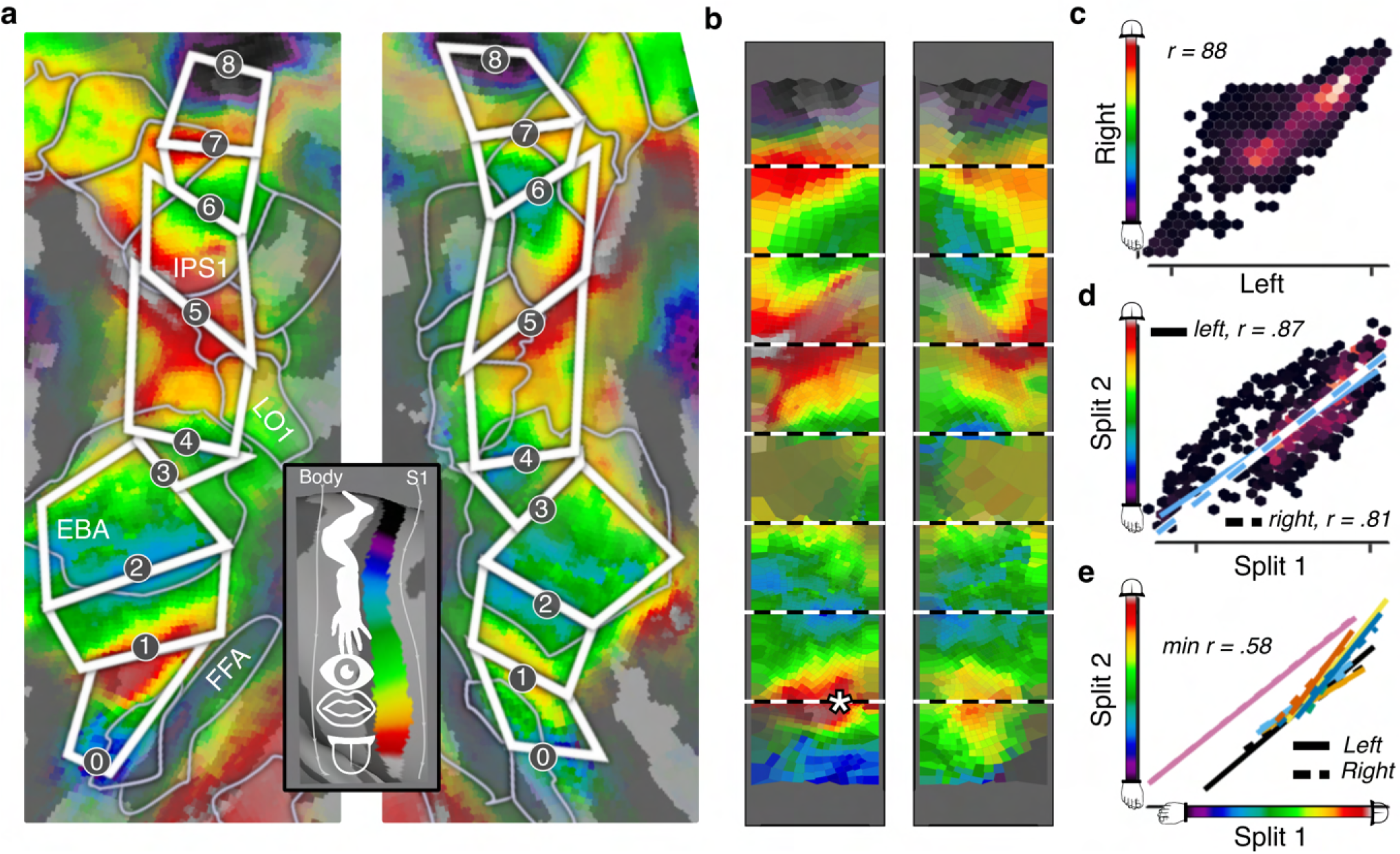
Dorsolateral visual cortex is tiled by at least 8 somatotopic maps during movie watching. **a.** Depicts the topography of connectivity with S1, as estimated by the connective field model) for the same cutout region as defined in Figure 3. The location of reversals are demarcated by white polygons to indicate gradient map boundaries. Vertices are thresholded to show only those with an out of set variance explained >10% and are transparency weighted between 10 and 40%. **b.** Voronoi plot of the polygons projected into a rectangular space. The asterisk demarcates the location of a strong left-hemisphere face representation that overlaps with the VWFA. **c.** Hexbin plot showing the agreement between data in the left and right hemisphere. **d.** Hexbin plot showing the agreement between independent subject splits, the least squares fit to the data is shown for each hemisphere. **e.** Least squares fits for the relationship between the data in each of the 16 maps across independent subject splits. The map is indicated by the color and the hemisphere is indicated by the linetype as shown by the legend.

To test the generalization performance of the somatotopic structure we employed cross-validation, which revealed that this somatotopic map predicted data from independent subjects with good accuracy (*r* = .84, p*<*10^−16^) (**Figure 4d**). We further validated the robustness of this body-map structure by a conservative permutation test that generates ‘surrogate’ somatotopic maps that maintain the autocorrelation structure of the empirical data (**see Methods**). This revealed that such out of sample agreement in somatotopic structure was rarely observed with surrogate instances (*p* = .026).

Notably, we observed the strongest hemispheric asymmetry in the most ventral map (max *p<*10^−3^ for pairwise comparisons of bootstrapped absolute hemispheric differences). The asymmetry of this map, which spans much of the region between EBA and FBA, was driven by stronger upper-limb/face representation in the left hemisphere (see asterisk in **Figure 4b**). Referencing this region against an atlas of occipito-temporal cortex,^26^ we observed that it overlaps with the visual word form area (VWFA)^27^ - a strongly left-hemisphere dominant region is functionally connected to S1 in both blind and non-blind individuals during manual exploration of tactile patterns.^28, 29^

### Somatotopic maps align with visual maps

Having established a somatotopic mode of organization in dorsolateral visual cortex, we next asked how it may be configured to connect to and recruit the computational machinery of vision. Since much of the visual system is retinotopically organized, one elementary way in which this may be implemented is via direct spatial alignment of the somatotopic and retinotopic reference frames. The most plausible form of this alignment would reflect ecological coincidences between visual field and bodily positions (e.g. feet often appear lower in the visual field) that optimize computation of environmental affordances (**Figure 5a**), and it could serve to facilitate processing related to interactions with the environment. To test this hypothesis, we performed a permutation-based searchlight analysis to detect local agreements between the somatotopic map and visual-field tuning estimates from the HCP retinotopy experiment (**Figure 5f**), **see Methods**). We detected evidence for this alignment mostly in dorsal regions that spanned V3B (*p<*10^−2.12^) and V3A (*p<*10^−2.02^) and more laterally in LO1 (*p<*10^−2.12^) and the superior portion of EBA (*p<*10^−1.90^), consistent with the EBA’s robust functional connectivity with the dorsal visuomotor stream^30^ and proposed role in action-related processing^19^ (**Figure 5b-f**).

**Figure 5.**
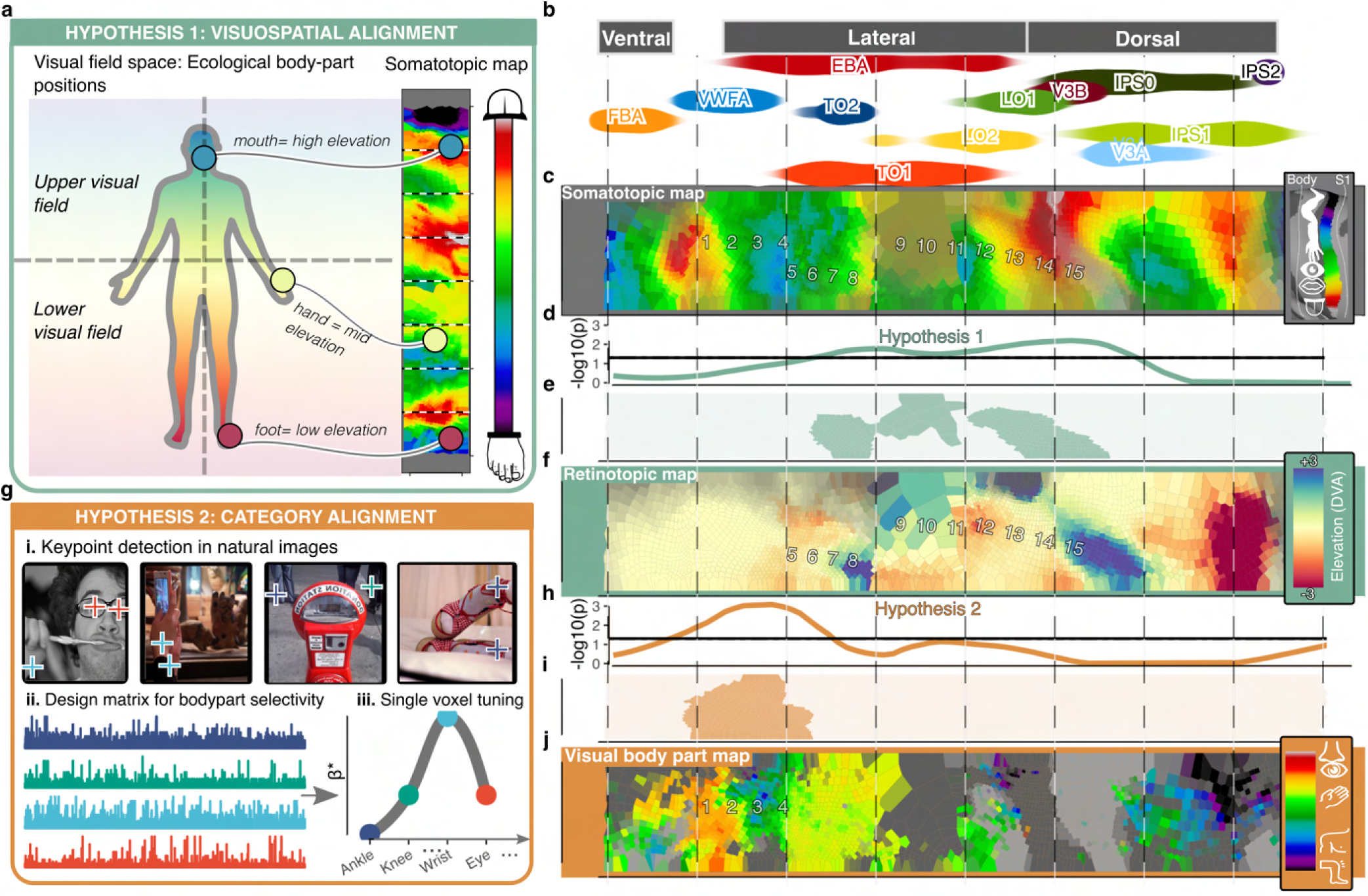
a. Schematic of the topographic alignment predicted by visuospatial alignment: Voxels tuned to upper visual field locations would be expected to coincide with parts of the somatotopic map corresponding to high-elevation body parts (and vice versa). **b.** Density plots depict the location of ROIs along the ordinate (roughly ventral-dorsal)) of the rectangular space defined in **4b**. **c.** Voronoi plot of same somatotopic map depicted in Figure 4b. Numbers are overlayed to aid visual comparison of corresponding points in ensuing figures. **d.** Evidence for alignment between the somatotopic map and retinotopic map shown in **f**, defined as maximum log10 p value of the chunkwise correlation. **e.** Depicts the geodesic chunks wherein evidence of alignment was detected. **f.** Depicts the retinotopic map, defined as vertical visual field position (y). **g.** Procedure for deriving map of visual bodypart selectivity. For NSD images, 17 anatomical keypoints were detected (4 shown to prevent overplotting) to compute predictions that reflect selectivity for each bodypart (see **Methods**). Image exemplars are those predicted to elicit the peak response for an eye, wrist, knee and ankle selective voxel respectively. **h.** Depicts the evidence for alignment between the somatotopic map and visual body part map shown in **j**, defined as the maximum log10 p value. **i.** Depicts the geodesic chunks for which evidence of alignment was detected. **j.** Depicts visual body part preference data generated from the procedure depicted in **g**. Colorbar depicts the position of bodily keypoints along this axis. All data in the right column are nearest-neighbor averaged across hemispheres. Searchlight analyses were performed separately on each hemisphere before averaging of outcomes was applied.

More ventrally, in the region overlapping with superior FBA, inferior EBA and the VWFA, we detected no evidence for the above-described spatial co-alignment between somatotopic and retinotopic tuning, which is narrowly biased towards the fovea. Although VWFA is not classically defined as a body-selective region to the same extent as FBA and EBA, it has been linked to responses to gestures of the hands and face.^29, 31^ Given the broader body-selective nature of this region, we reasoned that topographic co-alignment in this region may play out at a more categorical level, such that somatotopic structure predicts visual body-part selectivity. If this were the case, we should be able to predict, at the single-voxel level, visual selectivity to body parts from our somatotopic map.

To test this hypothesis, we leveraged the natural scenes dataset (NSD), a large 7T fMRI study of a total of around 300 scanning sessions wherein whole-brain responses were recorded from observers that viewed 10,000 natural images.^32^ We used a pose-estimation algorithm to code the presence of 17 anatomical ‘key points’ within each of the viewed images, ranging from lower body parts (e.g. ankle) to upper facial features (e.g. eyes). This visual body-part annotation formed the basis of a design matrix for a forward model of visual body-part selectivity, from which single-voxel visual body-part tuning could be derived **(Figure 5g**) **Methods**). This analysis provides a continuous metric of preferred visual body position preferences along a toe-tongue axis analogous to the somatotopic map (**Figure 5j**). The searchlight analysis revealed that the somatotopic map predicted visual body part preference from the superior portion of FBA into the ventral portion of EBA (*p<*10^−2.94^) and VWFA (*p<*10^−2.94^) **Figure 5h-j**). This finding of both ventral, visual-semantic alignment and dorsal, visuospatial alignment in EBA is consistent with its proposed role as an interface between dorsal ‘action’ and ventral ‘perception’ stream signals.^30^

## Discussion

A central question in sensory neuroscience is how inputs from vision and touch are combined to generate cohesive representations of the external world. Here we reveal a widespread mode of brain organization in which aligned, or ‘multiplexed’ topographic maps facilitate structured connections between vision and somatosensation. We developed a computational model that revealed somatotopic structure in dorsolateral visual cortex from movie watching data. Our analyses of somatotopic tuning in these regions found that this structure was predictive of both visual field locations and body part selectivity. Together, these results point towards a brain architecture with more extensive cross-modal overlap and interaction than traditionally assumed, as they indicate that the computational machinery classically attributed to the somatosensory system is also embedded within and aligned with that of the visual system. These topographically aligned, or multiplexed visual and bodily maps in humans are a likely brain substrate for non-afferent, internalized somatosensory representations of visual signals, and are a candidate human homologue of findings in mice where somato-motor responses dominate in visual cortex.^33^

Consistent with core predictions of embodied visual perception, our model-based quantifications of somatotopic and retinotopic connectivity revealed that dorsolateral visual cortical responses to naturalistic stimuli are best explained by selectivities in both modalities, as opposed to visual selectivities alone. The necessity of incorporating body-referenced processing into models of dorsolateral visual cortex aligns with a growing body of work indicating that its role extends beyond passive visual analysis, encompassing perceptual, semantic, and bodily functions optimized for behavioral interactions with the world.^19^

Consistent with a hypothesis of visuospatial alignment of somatosensory tuning, we found that body part preferences in dorsolateral visual cortex were predictive of visual field tuning. This alignment may be reinforced by shared developmental influences, as somatotopic and retinotopic maps are shaped trophically from birth, with the dorsal representation of the upper body and visual field and ventral representation of the lower body and visual field^17^, providing a roughly aligned sensory periphery optimized for efficient and systematic sampling and acting on the environment.^34, 35^ The explicit interweaving of touch and retinal coordinates we find in multiple dorsolateral regions may subserve efficient perception of environmental affordances and cohesive sense of self within space. Conversely, the imposition of a visuospatial reference frame on somatosensation likely contributes to the “rubber hand illusion” of spurious limb ownership, which has visuospatial dependence^36^ and to reduction of touch sensitivity in crossed arms,^37^ which is absent in the congenitally blind.^38^

Although aspects of this mapping may be stable and robust, others may be adaptable to new experiences. For instance, somatotopic organization in visual cortex reveals a body-referenced computational machinery shared between the visual and somatosensory systems that may explain substantial cross-modal plasticity: regions of dorsolateral visual cortex can support tactile discrimination tasks (e.g. Braille reading) in healthy subjects after less than a year of training.^28^ In addition to being prone to reorganization, the structured map-like organization betrays these regions’ flexibility. Due to their non-linear response properties,^39^ spatial selectivities in high-level visual cortex are characterized by substantial invariance to position and extent of stimuli: a flexibility that aids processes such as visual object recognition.^40^ It is plausible that this invariance principle extends to the level at which the modalities coincide - allowing for positional tolerance in registering common causes for visual and touch inputs. Extensions of our topographic connectivity framework may serve to chart these flexible interactions.

We also found that somatotopic connectivity in dorsolateral visual cortex predicted the existence and detailed tuning of a map of visual body-part selectivity which we map without reference to visual location. This congruence may underpin the high-level cross-modal modulations of body part representations that have been reported behaviorally: observing a body part can improve tactile localisation on that body part^41^ and visual judgements of body parts are enhanced when moving the corresponding body part.^42^ For such cross-modal modulations to be useful in the context of imitative visuomotor behaviors, they would require categorical visual representations of body parts that are robust to changes in size or viewpoint.

The convergence of somatosensory and visually referenced body part maps overlapped with the EBA, implicating this region in connecting felt and observed bodily sensations. This accords with causal evidence: electrical stimulation of EBA can lead to judgments of one’s own bodily sensations being biased towards observed bodily sensations.^43^ We also observed this body-part referenced tuning alignment in VWFA. Previous literature has shown that this region is activated by both visual word forms but also by communicative hand and face gestures.^31^ These characteristics together make this region a likely conduit between visual, body-related, and more abstract semantic computations important for language. It has been shown that electrical stimulation of sensorimotor regions supporting specific body movements (e.g. leg) can influence the processing of words semantically related to associated actions (e.g. kick). These observations support models of embodied language, which propose that lexical processing is supported by category-specific functional links between motor and language systems. Our findings highlight that this principle can be extended to additionally incorporate similar contributions from vision.^44^

Our findings also underscore the potency of naturalistic stimuli in revealing otherwise hidden multisensory interactions. In general, the brain exhibits stronger and more specific activations to naturalistic stimulation than to artificial stimuli.^45^ In the natural world, the brain must solve the problem of deriving meaning from fast multisensory stimuli embedded in extended contexts.^46^ Inference of a causal world model for effective behavior requires efficient fusion of sensory data across the senses, but the brain’s implementation of these processes has remained obscure. Topographic connectivity provides a tool to explore and track the neural processes responsible for cross-sensory fusion, both in parametric tasks and in the naturalistic contexts that confound traditional stimulus-referred analysis approaches. Here, we show that detailed information about the tuning of our sensory systems can be derived from neural responses during both rest and movie watching. This observation is promising for the study of conditions, including autism, that have aetiological roots in fundamental sensory computations, often manifesting as visual and tactile disturbances.^47^ The conventional approach to mapping these sensory systems via extended presentations of sparse stimuli presents intractable practical obstacles in populations prone to sensory hyper/hypo-sensitivity^48^ and co-morbid epilepsy.^49^ The stimulus-independent and task-free nature of our approach may therefore expand the viable playing field for revealing neural mechanisms underlying sensory dysfunction.

## Summary

We find that the reference frame of the human body permeates much of human cerebral cortex and is deeply embedded within the dorsolateral visual system. This integration casts dorsolateral visual cortex as a fundamentally multisensory part of the brain, whose multiplexed maps provide a rich canvas on which the brain can integrate the computational machinery of the different senses for various aspects of cognition. This appreciation of the role of dorsolateral visual cortex as a nexus between modalities is of interest in many conditions characterized by atypical interactions between vision and touch, such as autism spectrum disorder.^50^

## Acknowledgments

We thank Roni Maimon-Mor, Doug Saddy, Anastasia Christakou, Bhisma Chakrabarti, Paul Downing and Jesse Gomez for their comments on, and discussions about earlier versions of this manuscript. We would also like to thank the teams involved in the collection and curation of the Human Connectome Project Dataset and Natural Scenes Dataset.

## Methods

### Participants and Stimuli

fMRI Data were taken from the 174 participants of the HCP movie-watching dataset^1, 2^. The sample consisted of 104 females and 70 males (M age 29.3 years, SD = 3.3) born in Missouri, USA. 88.5% of the sample identified as ‘White’ (4.0% ‘Asian’, ‘Hawaiian or Other Pacific Island’, 6.3% ‘Black or African American’ 1.1% unreported). The English language comprehension ability of the sample (as assessed by age-adjusted NIH Picture Vocabulary Test^3^ scores) was above the national average of 100 (*M* = 110, *SD* = 15). Participants were scanned while watching short (ranging from 1 to 4.3 minutes in length) independent and Hollywood film clips that were concatenated into four movies of 11.9 - 13.7 minutes total length. Before each clip, and after the final clip was displayed, there were 20 second periods wherein there was no auditory stimulation and only the word ‘REST’ presented on the screen. There were 4 separate functional runs, wherein observers viewed each of the 4 separate movies. All 4 movies contained an identical 83 second ‘validation’ sequence at the end of the movie that was later removed to ensure independent stimulation in each cross-validation fold. Audio was scaled to ensure that no video clips were too loud or quiet across sessions and was delivered by Sensimetric earbuds that provide high-quality acoustic stimulus delivery while attenuating scanner noise. Participants also took part in one hour of resting state scans, also split into 4 runs of equal (15 min) length. Full details of the procedure and experimental setup are reported in the HCP S12000 release reference manual.^4^

### HCP data format and preparation

Ultra-high field fMRI (7T) data from the 174 subjects were used, sampled at 1.6 mm isotropic resolution and a rate of 1 Hz^1^. Data were preprocessed identically for movie-watching and resting state scans. For all analyses, the Fix independent component analysis-denoised time-course data, sampled to the 59,000 vertex-per-hemisphere areal feature-based cross-subject alignment method (MSMAll-^5^) surface format was used. These data are freely available from the HCP project website. The MSMAII method is optimised for aligning primary sensory cortices based on variations in myelin density and resting state connectivity maps^21^. Because of the unreliable relation between cortical folding patterns and functional boundaries, MSM method takes into account underlying cortical microarchitecture, such as myelin, which is known to match sensory brain function better than cortical folding patterns alone.^7^ Previous research has demonstrated that such an approach improves the cross-subject alignment of independent task fMRI datasets while at the same time decreasing the alignment of cortical folding patterns that do not correlate with cortical areal locations.^5^

We applied a high-pass filter to the timeseries data via a Savitzky Golay filter (3rd order, 210 seconds in length), which is a robust, flexible filter that allowed us to tailor our parameters to reduce the influence of low frequency components of the signal unrelated to the content of the experimental stimulation (e.g. drift, generic changes in basal metabolism). For each run, BOLD time series data were then converted to percent signal change.

For the purposes of cross-validation, we made training and test datasets from the full dataset. We removed the final 103 seconds of each functional run, which corresponded to the identical ‘validation’ sequence and final rest period at the end of each movie run. Our training dataset thus consisted of the concatenated data from the four functional runs with this final 103 seconds removed from each. The test dataset was created by concatenating the final 103 seconds from each run into a 412 second set of data.

All connective field models were fit on the individual subject data and for the movie-watching these models were also fit to the data of an across time-course averaged (‘HCP average’) participant. Split-half subject averages (*N*=87) were also created via a random 50% split of individual subject data. Split-half movie averages were created by creating separate datasets based on the first (movies 1 and 2) and second half (movies 3 and 4) of the movies.

### Connective Field Modeling

#### Model maps of V1 and S1 topography

Our analyses develop and extend the basic approach of connective field modeling, wherein responses throughout the brain are modeled as deriving from a ‘field’ of activity on the surface of a ‘source’ region - classically the primary visual cortex^8^ In turn, preferences for positions on the visual field can be estimated by referencing the estimated connective field V1 positions against the retinotopic map of V1 (**Figure 1a-c**). Here, we extend this approach by simultaneously modeling brain responses as resulting from connective fields on both the V1 and S1 surface. As such, this minimally requires us to estimate both a V1 and S1 source region and their underlying retinotopic and somatotopic map.

To define these V1 and S1 source regions, we first made sub-surfaces (one for each hemisphere) from the full cortical mesh, containing the vertices corresponding to V1 and S1 as defined by coordinates from a multi-modal parcellation of the HCP data for regions V1 and 3b.^21^

To provide a model retinotopic map for V1, we used data from a ‘retinotopic prior’ that defines subject-averaged parameters of preferred visual field position (eccentricity, polar angle) estimated from a population receptive field (pRF) analysis of the HCP data.^9^ Thus every vertex in the V1 was associated with an eccentricity and polar angle value that defined its preference for the corresponding position in the visual field. Using this data, the V1 subsurface was then additionally curtailed to only include vertices that fell within the region stimulated by the movie display (within 8 DVA from the fovea).

The known topographic organization of S1 is a somatotopic map, which is an approximately dorsomedially to ventrolaterally oriented gradient of sensitivity that runs from sensitivity to lower limbs to the upper limbs and face.^12^ To first provide a continuous coordinate space for this somatotopic gradient, for each vertex in S1, we calculated the geodesic distance from the vertices at the most dorso-medial edge of the S1 subsurface. Thus every vertex in S1 was associated with a value reflecting its distance from the dorso-medial edge of S1.

To explicitly relate these S1 coordinates to body parts, we leveraged data from an independent, publicly available motor mapping experiment, collected at 3T, where 62 participants performed movements with 12 discrete body parts ranging from toe to tongue.^15^ These subject-wise beta weights for each bodypart were nearest-neighbor resampled into the same 59k vertex per hemisphere space as the HCP data. A second-level random effects GLM analysis was then conducted to reveal the S1 positions corresponding to the peak of the group-level *t* statistic for each bodypart.

#### Design matrix

We first summarized the V1 and S1 sub-surfaces as a finite set of spatial profiles by deriving eigenfunctions of their Laplace-Beltrami operator (LBOEs). This decomposition, which has been referred to as recovering the ‘shape DNA’^12^ of a manifold, yields a finite family of real-valued functions that are intrinsic to the surface shape, orthogonal and ordered according to spatial scale^13^ (**see Extended Data Figure E1a**). In principle therefore, one can approximate any arbitrary spatial pattern on the surface (i.e. a connective field) via a linear combination of LBOEs.

To validate this approach, and determine the number of LBOEs to use in our analysis, we conducted pilot analyses where we attempted to predict target Gaussian connective fields of varying sizes from linear combinations of LBOEs. These analyses indicated that for both V1 and S1, 200 LBOEs were sufficient to adequately predict connective fields with a sampling extent of 2°mm, which approximates the lower bound of the known sampling extent of extrastriate cortex from V1 (i.e. in V2).^8^ Reconstruction performance was at near-ceiling levels for sampling extents of 4mm and above (**see Extended Data Figure E2,Extended Data Figure E3**) and increasing the number of LBOEs from 200 led to relatively trivial increases in reconstruction performance, with large increases in computation time. Furthermore, visualizing the performance of predictions for target connective fields centered on each V1 and S1 vertex revealed no systematic spatial inhomogeneities in the performance of models based on 200 LBOEs (**Extended Data Figure E3).** As such we opted to use 200 LBOEs per subsurface in our connective field modeling.

We next generated model timecourses for our design matrix via the dot product of the timecourse data in each subsurface and each of the 200 corresponding LBOEs. Each model timecourse therefore reflects the sum of the timeseries data within the subsurface, weighted by an orthogonal spatial profile (**see Extended Data Figure E1b**). The model timecourses were then z-scored over time and stacked to form a design matrix for model fitting. Thus, there were 800 regressors in our design matrix: 400 from V1 and 400 from S1 (200 per hemisphere) which were used to explain the resting-state and movie-watching data.

#### Model fitting

All model fitting was conducted in python, exploiting the routines implemented by the ‘Himalaya’ package.^14^ In ordinary least-squares (OLS) regression, one estimates weights *b*, such that data *y is* approximated by a linear combination of regressors *Xb* (**see Extended Data Figure E1c**). Here, we employed banded ridge regression,^14^ which belongs to a family of ‘regularized’ regression techniques that estimate a regularization parameter *λ* to improve the generalization performance of OLS regression.^15^ Banded ridge regression expands on these techniques by estimating a separate *λ* for separate feature spaces *i* of the design matrix *X* - thereby optimizing regularization strengths independently for each feature space (**see Extended Data Figure E1d**). Banded ridge regression therefore respects the fact that different feature spaces in the design matrix may differ in covariance structure, number of features and prediction performance - entailing different optimal regularization.

In the present case, our two feature spaces consisted of the visual and somatosensory modalities, or equivalently, the 400 V1 and S1 model timecourses (X*_v1_*, X*_s1_*) described in the previous section. Thus, to model brain activity of a particular voxel, banded-ridge regression computes the weights *b* _i_*, as defined below:

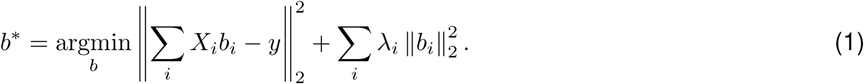

Similarly to un-banded ridge regression, the ridge weights *b*_i_*are estimated from the training data and the hyperparameters *λ* _i_ are learned via cross validation. In the present case, our training data consisted of four runs of functional data wherein participants watched an independent movie. This natural organization of the data allowed us to use a leave one movie out cross-validation strategy to estimate *λ _i_*.

#### Connective field estimation

For a given voxel, the coefficients *b* _i_*estimated by the banded ridge regression model can be interpreted as the cross-validated ‘importance’ of each model timecourse in the design matrix in explaining its response throughout the movie-watching. By extension, since each model timecourse derives from an orthogonal spatial profile on the surface of V1 or S1, this means that *b*_i_* also implicitly estimates the ‘importance’ of each of the underlying spatial profiles. Accordingly, for any given voxel, the dot product of its estimated *b* _i_* and the corresponding spatial profiles *s_i_*reveals a spatial map of the importance of each vertex on S1 and V1 in explaining the voxels response - or equivalently - it estimates its visual and somatosensory ‘connective field’ (**see Extended Data Figure E1e**).

Notably, this method of estimating the connective field from a finite set of weighted spatial profiles differs from the ‘classic’ connective field estimation procedure as it removes the constraint that the connective field profile is a Gaussian defined by a center (*V0*) and extent *(σ)* - and can estimate an infinite number of arbitrary spatial profiles of a smoothness depending on the amount of LBOEs entered into the analysis.

#### Connectivity-derived retinotopic and somatotopic mapping

With connective field profiles estimated for each vertex, preferred visual field and body positions were estimated by taking the dot product of each S1 and V1 connective field and the corresponding somatotopic and retinotopic map (see **Model Maps of V1 and S1 Topography**) and then dividing by the sum of the connective fields (**see Extended Data Figure E1f**). Since the values of the connective field profile represent the importance of each location on the source region in explaining a given voxels response, this is akin to a weighted averaging, whereby the retinotopic/somatotopic maps are averaged in a manner weighted by the predictive performance at each location.

#### Model performance metrics

The joint specification of our model with multiple feature spaces also allows us to disentangle the contribution of each feature space (sensory modality) to overall prediction performance. Specifically, variance decomposition via the product measure, allows the computation of independent *R*^2^ scores per feature space that sum to the total *R*^2^ of the joint model.

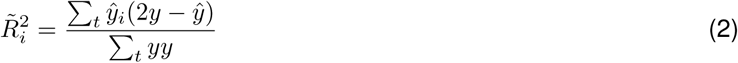

Where 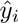 is the sub-prediction computed on feature space *X_i_* alone, using the weights *b* _i_*of the joint model. To evaluate out-of-sample performance of the model, the parameters estimated from the training data were then used to predict the test data and the variance explained from the V1 and S1 feature spaces was evaluated using the above formula.

### ROI definitions

#### Classically Somatotopic ROIs

To define somatotopic regions of interest, we leveraged a previously defined, gross anatomical parcellation of parietal, medial, insular and frontal zones that have been found to contain robust homuncular somatotopic gradients, or ‘creatures’ of the somatosensory system.^12^ These ‘creatures’ were themselves defined by a combination of regions in the multi-modal parcellation defined by Glasser and colleagues,^21^ whose voxel-averaged response to tactile stimulation was significantly above zero. Further details of the exact regions of the Glasser parcellation that correspond to each ROI can be found in the Saadon-Grosman paper.

#### Classically Visual ROIs

To define visual regions of interest, we leveraged a pre-existing probabilistic atlas of 25 retinotopic visual regions provided by Wang and colleagues.^20^ To this parcellation, we added the regions FFA, FBA, EBA and PPA, which were defined by functional localizer (floc) data taken from the Natural Scenes Dataset (NSD).^32^ Specifically, the subject-averaged t statistics for the faces/bodies v all other stimulus categories contrast were thresholded at the *α<*.05 level and ROIS were hand-drawn using pycortex. Note therefore, that we opted to not use the pre-drawn definitions packaged with the NSD dataset, which were defined according to a liberal *t >*0 thresholding. Our definition of the EBA overlaps with the Wang regions LO2, TO1 and TO2, hence only the EBA is displayed on cortical flatmaps to avoid overplotting. The relationship between these regions is shown in (**Extended Data Figure E7a**).

### Statistical testing

#### Topographic connectivity scores

To generate a measure of topographic connectivity, the out-of-set *R*^2^ for visual and somatosensory connective field predictions were corrected for the *R*^2^ of a non-topographic ‘null’ model, whose predictions were generated via the mean V1 and S1 timecourses respectively. This means that although the corrected values are no longer interpretable as variance explained, they reflect the superiority of the generalization performance of a spatial connective field model relative to a non-spatial model. This correction thus conservatively assesses the presence of true topographic connectivity by referencing against an explicit null model. One sample t tests were conducted to compare these corrected scores against zero and reported *p* values are two sided. To provide estimates of the magnitude of topographic connectivity, we computed the effect size Cohen’s *dz* via the formula:

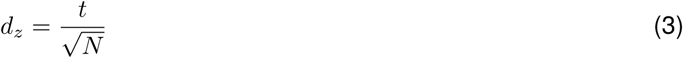

Within subjects differences in topographic connectivity scores were analyzed via repeated measures ANOVA, implemented in the “afex” package in the R programming language.^18^ In modeling pairwise differences in topographic connectivity scores between ROIS, we performed holm-bonferroni correction of p values to account for the number of tests conducted. This was implemented via the routines implemented in the “emmeans” R package.^19^

#### Cortical coverage of somatotopic connectivity and relation to functional localiser

The extent of cortical coverage was assessed via a standard bootstrapping procedure. Individual subject topographic connectivity scores were resampled with replacement 10,000 times to generate resampled datasets with random samples of participants. For each resampled dataset, we quantified the percentage of voxels within each somatosensory ROI with topographic connectivity scores significantly greater than zero. In this calculation, note that the source region of the analysis (S1) is excluded. Reported confidence intervals were obtained from the quantiles (2.5% and 97.5%) of the resulting distribution of % coverage estimates. The same bootstrapping approach was applied to assess the correlation of somatotopic connectivity scores with functional localiser data from the NSD dataset, for which we used the same group-level t statistics described above. Using the correlation between somatotopic connectivity scores and the ‘body v all other categories’ *t* statistics as a reference, we subtracted the correlation with the corresponding place, face and object *t* statistics across 10,000 bootstrapped samples. The resulting distributions of correlation differences were used to compute *p* values.

#### Robustness of extrastriate somatotopic maps: permutation test

To evaluate the robustness of extrastriate somatotopic maps, we tested the null hypothesis that out of set somatotopic maps are predicted by maps generated from randomized connective fields, but with preserved autocorrelation structure. Referencing the out of set prediction of empirical maps against surrogate instances is important, since the spatial autocorrelation inherent in brain data implies that spatially proximal measurements are likely to be similar, regardless of how they were derived.^20^ As such, the statistical significance of agreement between maps is likely to be inflated and violate independence assumptions. Thus, rather than relying on the statistical significance of these empirical agreement statistics alone, they require benchmarking against surrogate data that quantifies the statistical expectations under a null hypothesis.

To this end, we first used the somatotopic map estimated for the cutout region in **Figure 4a** from one split-half of subjects to predict the corresponding data from the other split-half. The resulting *R*^2^ value was retained as an empirical test statistic. To generate a null distribution of these statistics, we generated 10,000 ‘surrogate’ homuncular maps from each split-half of subject data. These were generated by taking the estimated *b\** corresponding to each of the LBOEs of the S1 subsurface, randomizing their sign and recomputing the S1 connective field. This manipulation generates randomised connective fields that preserve the same amplitude spectra (distribution of energy across frequency) as the empirical data. For each surrogate dataset, we then computed its performance in predicting the out of set empirical somatotopic map and retained the proportion of these statistics that exceeded the empirical value as a *p* value. This process was repeated for the second subject split and the resulting *p* values were summed to obtain a final measure of the probability of obtaining the empirical statistic under the null hypothesis.

#### Visual body-part selectivity estimates

To estimate a map of visual body-part selectivity, we leveraged data from the Natural Scenes Dataset (NSD), also collected at 7T.^32^ Specifically, we used the denoised single trial beta-estimates from the final 12 runs of data of all 8 subjects. This corresponded to 9000 functional volumes, each of which included voxel-wise estimates of the response to an image from the common objects in context (COCO) dataset.^21^ Using Connectome Workbench commands,^22^ the cortex-wide single-trial beta-estimates were nearest-neighbor resampled from fsaverage space to the same 59k vertex-per-hemisphere surface format as the HCP data.

We next leveraged a corpus dataset of estimated body part keypoints within each image of the COCO dataset, which were generated by Openpose,^23^ a convolutional neural network based pose-estimation toolkit. Openpose detects the location of 17 different keypoints (nose, left eye, right eye, left ear, right ear, left shoulder, right shoulder, left elbow, right elbow, left wrist, right wrist, left hip, right hip, left knee, right knee, left ankle, right ankle). Thus, each image in the COCO dataset is associated with 17 binary variables that codes the presence of each keypoint for every human entity within the image. Critically, the subset of COCO images used in the NSD dataset were spatially cropped relative to the original versions for which the keypoints were computed. We therefore re-coded keypoints that were coded as present in the original COCO images but resided outside of the NSD crop-box as being 0 (absent). For each image and bodypart, we calculated the average of the binary variable for each keypoint across entities to give an estimate of the frequency with which the bodypart was present within entities within the image.

Next, we converted these scores into a regressor that explicitly coded selective responses for each bodypart. For each bodypart, we calculated a bodypart selectivity score defined as follows:

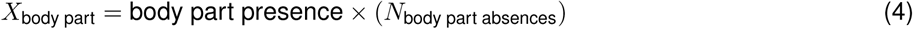

In this calculation, for example, an image with the presence of an ankle and the absence of all other body parts generates the highest score for the ankle selectivity regressor. Conversely, the score for an image will be implicitly penalized as a function of the number of other (non-ankle) body parts that are visible. These regressors for each body part were stacked into a design matrix. This formed the basis of a forward model of visual body selectivity, which was estimated by ridge regression. The model was trained on 10 of the runs of data via k-fold cross-validation and was tested on the final 2. For each voxel, body part preference was defined as the dot product of the 12 resulting beta weights and their ordinal position in the S1 homunculus (ankle to nose) providing a continuous map of visual body part selectivity along a similar toe-tongue axis as the somatotopy data.

#### Searchlight analyses: permutation test

Within the cutout region in **Figure 4** we defined geodesic ‘chunks’ of cortex that were centered at each vertex with a radius of 8mm. To generate empirical statistics, within each one of these chunks we computed the correlation between the somatotopic map shown in **Figure 4** and the target data (retinotopic map or visual body-part selectivity map). To generate a null distribution of these statistics, we leveraged the same 10,000 phase-scrambled ‘surrogate’ homuncular maps (see **Extrastriate homuncular maps: permutation test**) and for each one we calculated the corresponding correlations with the target data in each chunk. This distribution of local chunk-wise correlations obtained across all surrogate somatotopic maps served as our null distribution of local correlations. *p* values for the correlations within each empirical chunk were obtained as the probability of obtaining such extreme statistics from this null distribution. Note that for the analysis of the retinotopy data, we excluded chunks wherein the range of estimated visual field positions was less than 1DVA and so had little retinotopic variation. The resulting *p* values were then projected into the spatial locations of their corresponding chunk and the data were then averaged across hemispheres to produce the data in **Figure 5e,i**. Note that since the chunks contained partially overlapping data, the data in **Figure 5** show the lowest *p* value obtained at each vertex location.

## Extended Data

**Figure E1.**
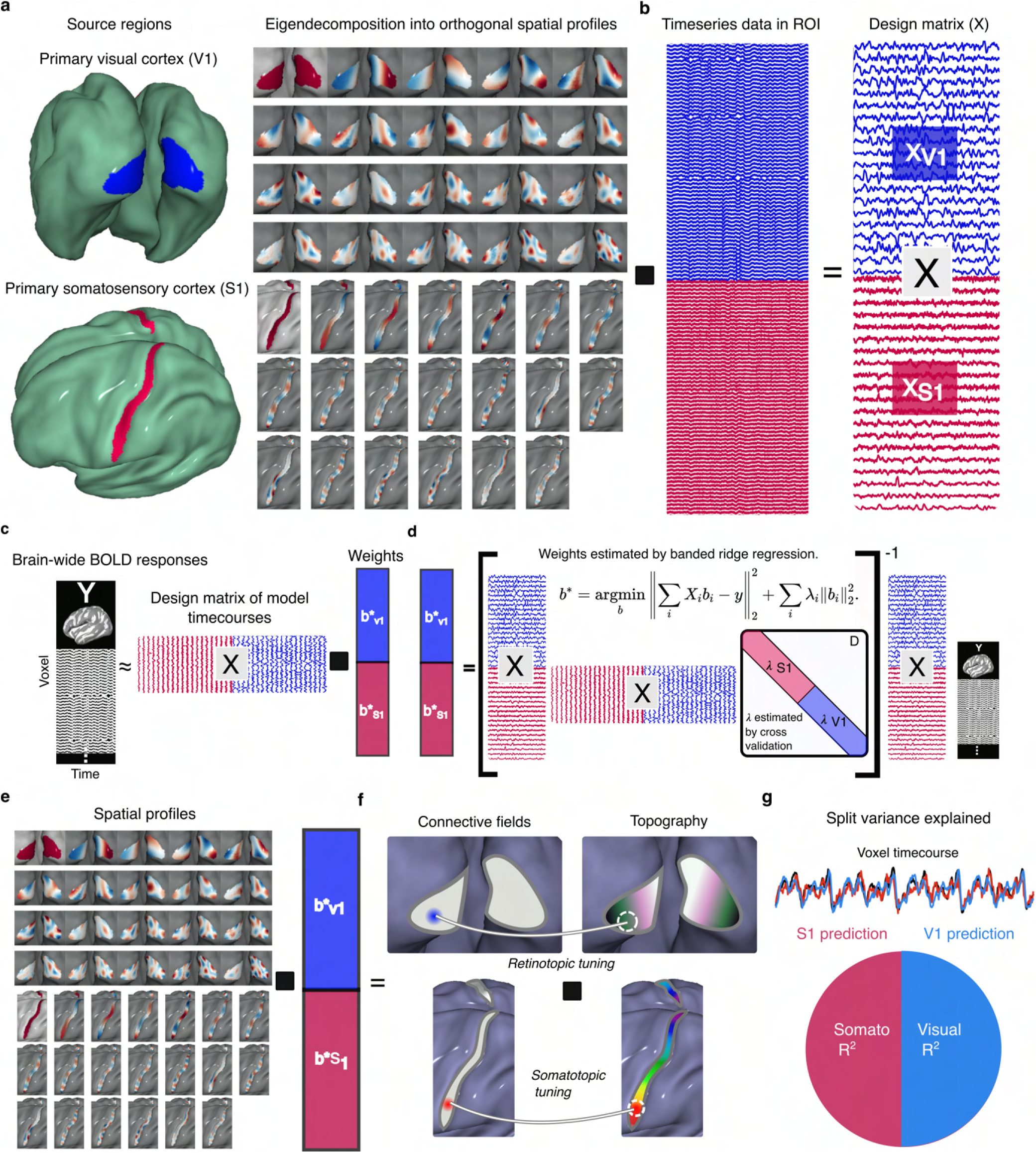
Illustration of Multisensory Connective Field Modeling Procedures. **a.** Initial source regions are defined from the vertices corresponding to V1 and S1, from which 200 LBOEs are defined. **b.** The dot product of these spatial profiles and the BOLD timecourses for each vertex within each subsurface produces a banded design matrix for a GLM. **c.** For each voxel in the brain, its BOLD timecourse is estimated as a linear combination of the model timecourses in the design matrix. **d.** Parameters of the GLM are estimated by banded ridge-regression, employing a leave one run out cross-validation scheme. **e.** With ridge beta weights (b*) estimated, the connective field for V1 and S1 is estimated as the dot product of the b* and the corresponding LBOEs. **f.** The retinotopic and somatotopic tuning of each voxel is then estimated as the weighted average of its estimated connective field profiles and the topographic map of each source region. **g.** The dual, banded nature of the design matrix allows the variance explained to be partitioned into the split proportion explained by V1 and S1 connective fields. These split R^2^ values that sum to the total variance explained. **See Methods for more details.**

**Figure E2.**
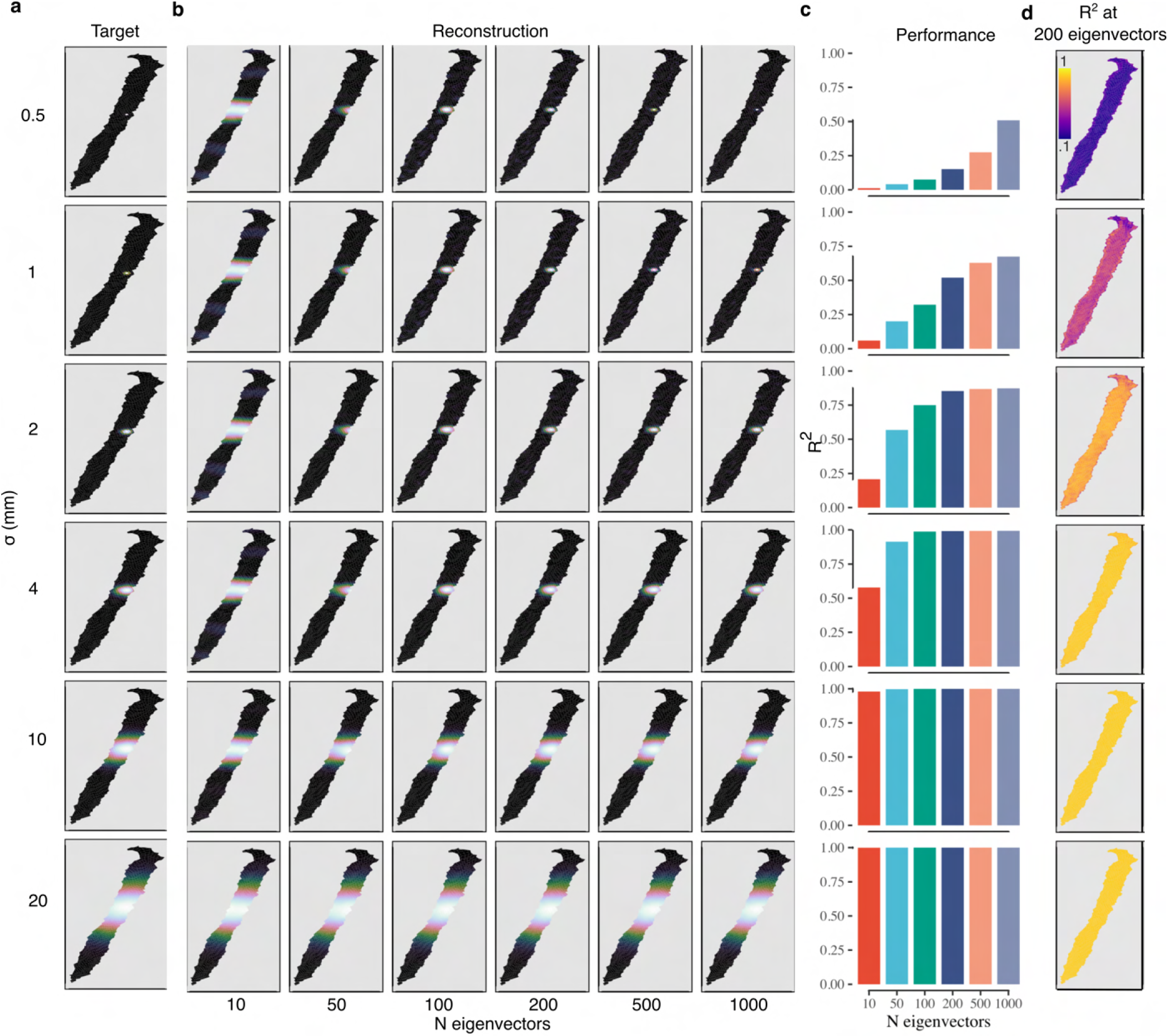
Validation of Connective Field Estimation: S1. **a.** Depicts a target connective field on the surface of S1, defined by a Gaussian function with a standard deviation (σ) defined in mm. **b.** The predicted connective field as a function of the number of S1 LBOEs in the design matrix. **c.** The agreement (defined by R^2^) between the predicted and target connective fields as a function of σ and number of LBOEs. **d.** For each vertex in S1, depicts the variance explained for a model with 200 LBOEs.

**Figure E3.**
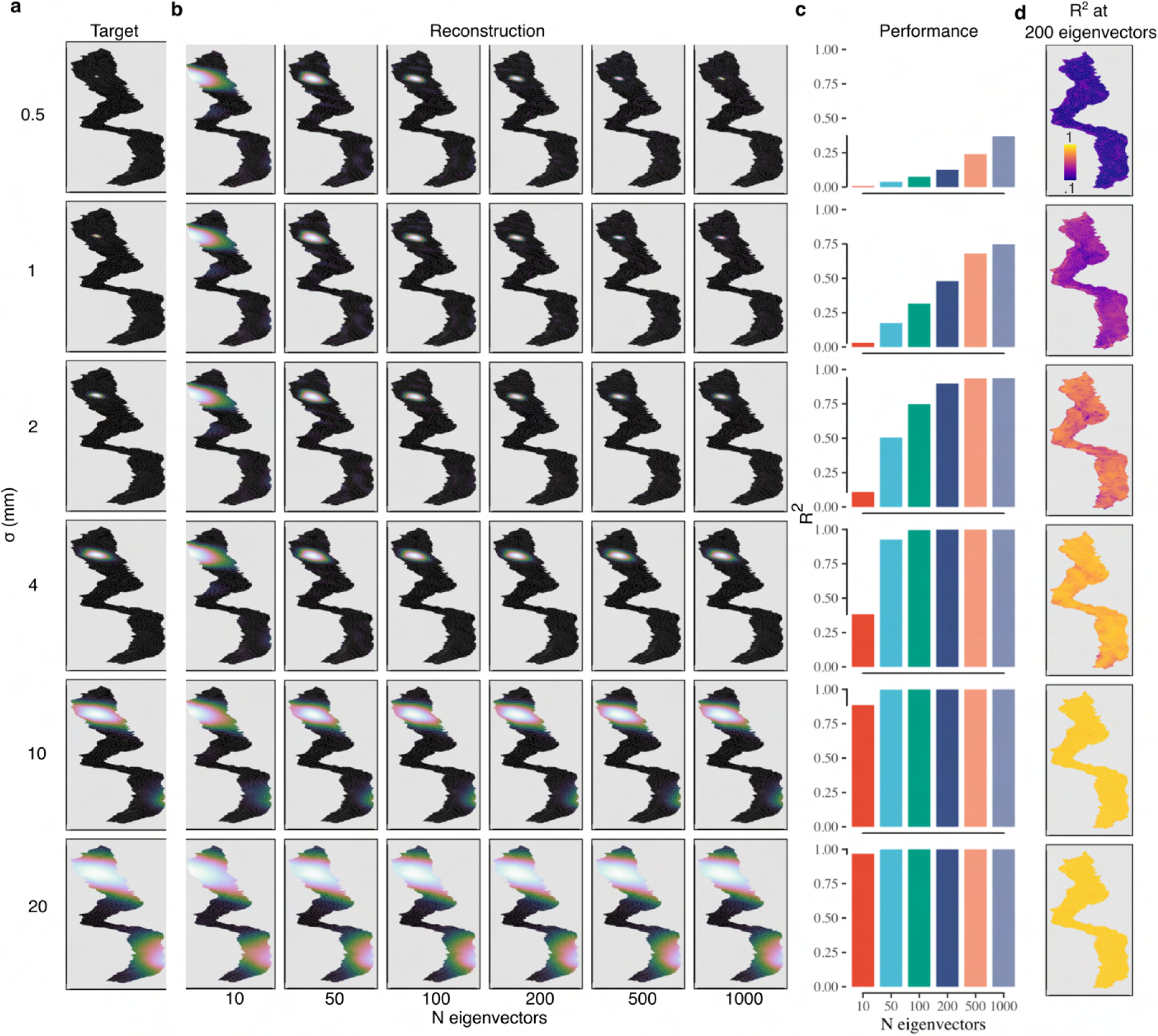
Validation of Connective Field Estimation: V1. **a.** Depicts a target connective field on the surface of V1, defined by a Gaussian function with a given standard deviation (σ). **b.** The predicted connective field as a function of the number of V1 LBOEs in the design matrix. **c.** The agreement (defined by R^2^) between the predicted and target connective fields as a function of σ and number of LBOEs. **d.** For each vertex in V1, depicts the variance explained for a model with 200 LBOEs.

**Figure E4.**
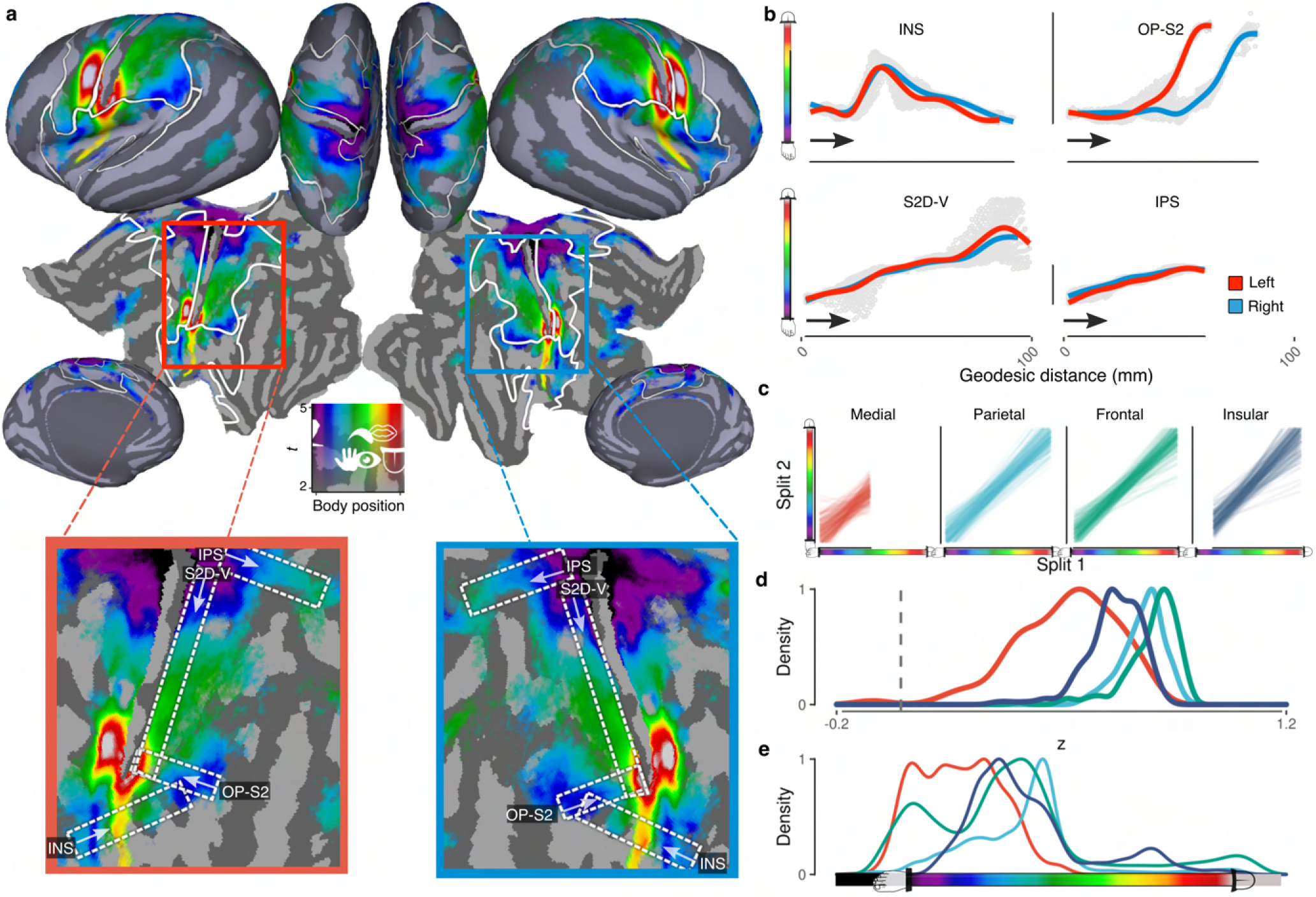
Structure and reliability of resting state somatotopy. **a.** Shows preferred body position (colors) derived from connective field modeling of resting state data. **b.** For each hemisphere (colors) shows the relationship between geodesic distance and body position along each of the gradients depicted in the lower panel of **a. c.** Lines represent least-squares fits for the relationship between somatotopic maps of independent subject splits. Different panels/colors show the data for different somatotopic rois, which correspond to the solid white lines in **a**. **d.** Density plot depicting the distribution of Fisher-Z transformed correlations for the data shown in **c. e.** Density plot depicting the distribution of body-part selectivity for each somatotopic roi. Colors in **d-e** correspond to the roi labels in **c**.

**Figure E5.**
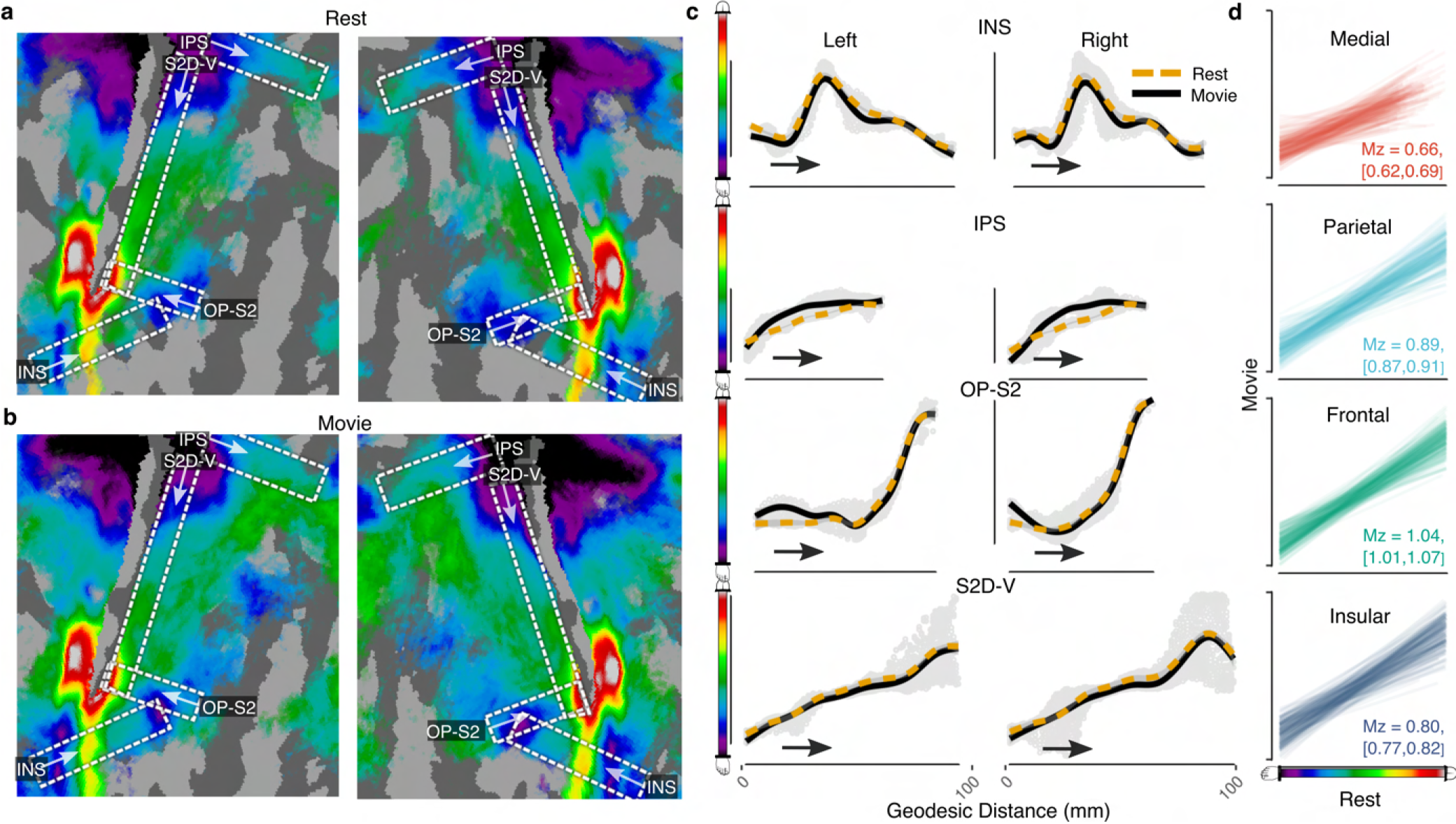
Comparison of somatotopy derived from rest and movie-watching. **a.** Shows preferred body position (colors) derived from connective field modeling of resting state data (same data as **Extended Data Figure E4a**). **b.** For comparison, the same outcomes are plotted from the movie-watching data. **c.** For each hemisphere and task (rest, movie) depicts the relationship between geodesic distance and body position along each of the gradients depicted **a/b. d.** Lines represent least-squares fits for the relationship between the somatotopic maps of rest and movie watching data from each subject. The text in each panel indicates the mean Fisher-z transformed correlation and 95% CI.

**Figure E6.**
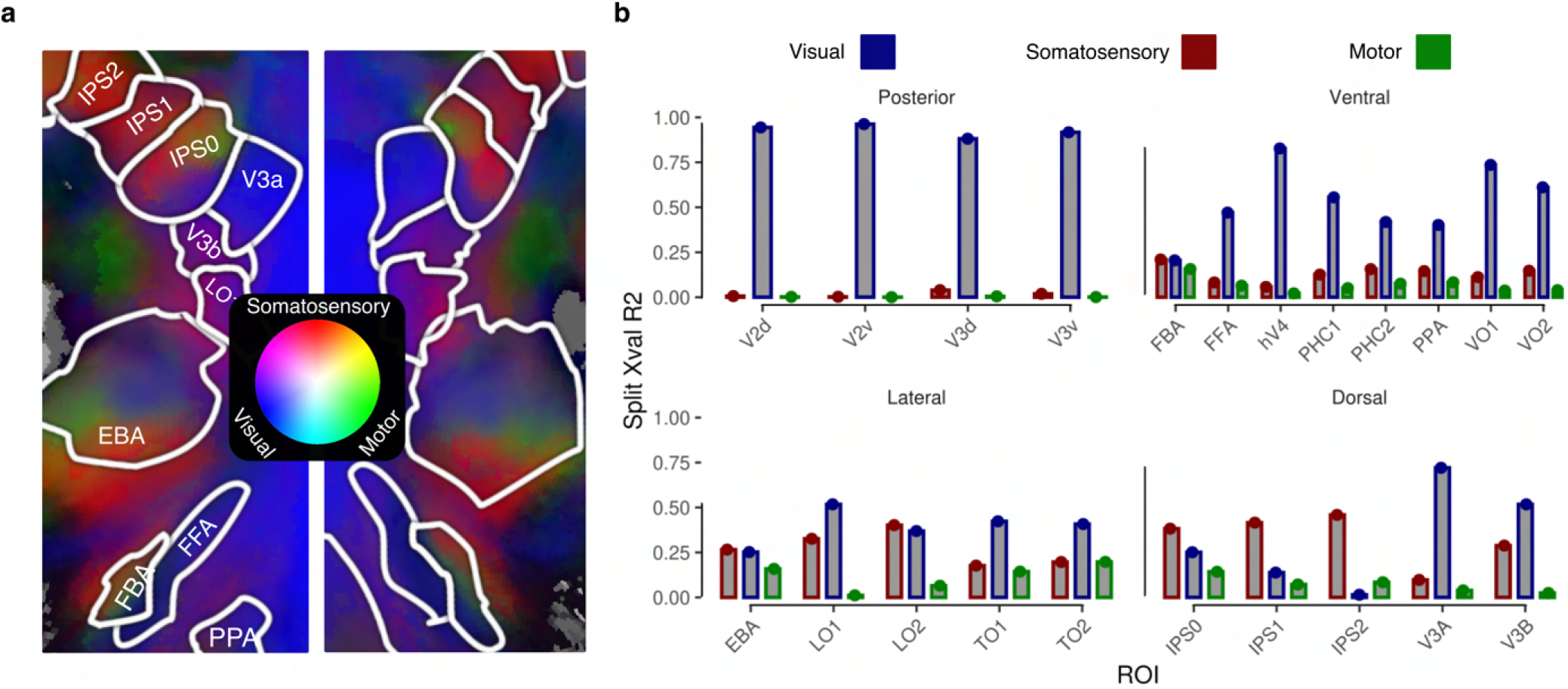
Outcomes from triple source connective field modeling with M1. RGB colormap Depicts the cross-validated split variance explained by triple-source connective field modeling from S1(somatosensory), M1 (motor) and V1 (visual). **b.** Bar charts depict visual cortical region-based average of S1, M1 and V1-derived cross-validated variance explained for the respective connective field model. In all regions of visual cortex, responses are better explained by connective fields deriving from S1 than from M1.

**Figure E7.**
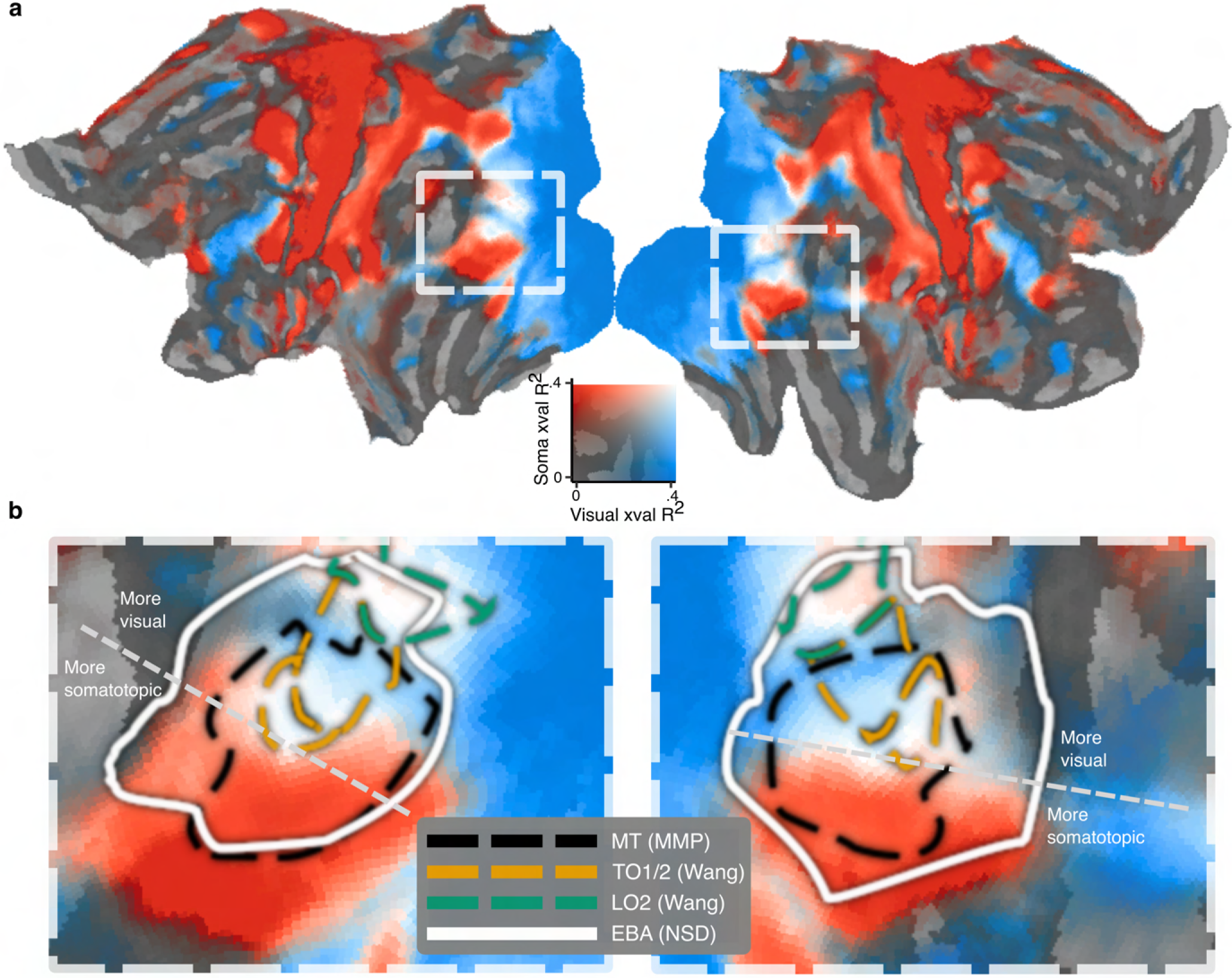
Organisation of EBA and overlapping regions in relation to multimodal topographic connectivity. **a.** Connective field strength from each of the source regions (V1 - blue, S1 - red) is quantified by their separate cross-validated explained variance. The dashed rectangle depicts an area of lateral visual cortex that encompasses the EBA. **b.** Cutout depicts zoomed-in version of the data within the rectangle defined in **a.** The legend depicts ROI definitions for EBA, retinotopic regions LO2, TO1-2 and the human MT complex. An additional dotted white line is drawn to demarcate a modality shift to dominant somatotopic connectivity, which aligns well with the border of TO2.

## References

1. Buccino, G. et al. Action observation activates premotor and parietal areas in a somatotopic manner: An fMRI study. Eur J Neurosci 13, 400–404 (2001).

2. Kuehn, E., Haggard, P., Villringer, A., Pleger, B. & Sereno, M. I. Visually-Driven Maps in Area 3b. J. Neurosci. 38, 1295–1310 (2018).

3. Astafiev, S. V., Stanley, C. M., Shulman, G. L. & Corbetta, M. Extrastriate body area in human occipital cortex responds to the performance of motor actions. Nat Neurosci 7, 542–548 (2004).

4. Orlov, T., Makin, T. R. & Zohary, E. Topographic Representation of the Human Body in the Occipitotemporal Cortex. Neuron 68, 586–600 (2010).

5. Keysers, C., Kaas, J. H. & Gazzola, V. Somatosensation in social perception. Nat Rev Neurosci 11, 417–428 (2010).

6. Gazzola, V., Aziz-Zadeh, L. & Keysers, C. Empathy and the Somatotopic Auditory Mirror System in Humans. Current Biology 16, 1824–1829 (2006).

7. Gonzalez-Castillo, J., Kam, J. W. Y., Hoy, C. W. & Bandettini, P. A. How to Interpret Resting-State fMRI: Ask Your Participants. J. Neurosci. 41, 1130–1141 (2021).

8. Delamillieure, P. et al. The resting state questionnaire: An introspective questionnaire for evaluation of inner experience during the conscious resting state. Brain Research Bulletin 81, 565–573 (2010).

9. Proffitt, D. R. An Embodied Approach to Perception: By What Units Are Visual Perceptions Scaled? Perspect Psychol Sci 8, 474–483 (2013).

10. Groen, I. I. A., Dekker, T. M., Knapen, T. & Silson, E. H. Visuospatial coding as ubiquitous scaffolding for human cognition. Trends in Cognitive Sciences 26, 81–96 (2022).

11. Wandell, B. A. & Winawer, J. Imaging retinotopic maps in the human brain. Vision Res. 51, 718–737 (2011).

12. Saadon-Grosman, N., Loewenstein, Y. & Arzy, S. The ‘creatures’ of the human cortical somatosensory system. Brain Commun 2, fcaa003 (2020).

13. Knapen, T. Topographic connectivity reveals task-dependent retinotopic processing throughout the human brain. Proceedings of the National Academy of Sciences 118, e2017032118 (2021).

14. Zeharia, N., Hertz, U., Flash, T. & Amedi, A. New Whole-Body Sensory-Motor Gradients Revealed Using Phase-Locked Analysis and Verified Using Multivoxel Pattern Analysis and Functional Connectivity. Journal of Neuroscience 35, 2845–2859 (2015).

15. Ma, S. et al. An fMRI dataset for whole-body somatotopic mapping in humans. Sci Data 9, 515 (2022).

16. Schneider, R. & Gautier, J. C. Leg weakness due to stroke. Site of lesions, weakness patterns and causes. Brain 117 (**Pt 2**), 347–354 (1994).

17. Flanagan, J. G. Neural map specification by gradients. Curr Opin Neurobiol 16, 59–66 (2006).

18. Pitcher, D. & Ungerleider, L. G. Evidence for a Third Visual Pathway Specialized for Social Perception. Trends Cogn Sci 25, 100–110 (2021).

19. Lingnau, A. & Downing, P. E. The lateral occipitotemporal cortex in action. Trends in Cognitive Sciences 19, 268–277 (2015).

20. Wang, L., Mruczek, R. E., Arcaro, M. J. & Kastner, S. Probabilistic Maps of Visual Topography in Human Cortex. Cereb Cortex 25, 3911–3931 (2015).

21. Glasser, M. F. et al. A multi-modal parcellation of human cerebral cortex. Nature 536, 171–178 (2016).

22. Peelen, M. V. & Downing, P. E. Is the extrastriate body area involved in motor actions? Nat Neurosci 8, 125–125 (2005).

23. Gallese, V., Keysers, C. & Rizzolatti, G. A unifying view of the basis of social cognition. Trends in Cognitive Sciences 8, 396–403 (2004).

24. Peelen, M. V. & Downing, P. E. The neural basis of visual body perception. Nat Rev Neurosci 8, 636–648 (2007).

25. Heimann, K., Umilta, M. A., Guerra, M. & Gallese, V. Moving mirrors: A high-density EEG study investigating the effect of camera movements on motor cortex activation during action observation. J Cogn Neurosci 26, 2087–2101 (2014).

26. Rosenke, M., van Hoof, R., van den Hurk, J., Grill-Spector, K. & Goebel, R. A Probabilistic Functional Atlas of Human Occipito-Temporal Visual Cortex. Cerebral Cortex 31, 603–619 (2021).

27. Dehaene, S. & Cohen, L. The unique role of the visual word form area in reading. Trends Cogn Sci 15, 254–262 (2011).

28. Siuda-Krzywicka, K. et al. Massive cortical reorganization in sighted Braille readers. eLife 5, e10762 (2016).

29. Debska, A., Wojcik, M., Chyl, K., Dziegiel-Fivet, G. & Jednorog, K. Beyond the Visual Word Form Area – a cognitive characterization of the left ventral occipitotemporal cortex. Frontiers in Human Neuroscience 17 (2023).

30. Zimmermann, M., Mars, R. B., de Lange, F. P., Toni, I. & Verhagen, L. Is the extrastriate body area part of the dorsal visuomotor stream? Brain Struct Funct 223, 31–46 (2018).

31. Xu, J., Gannon, P. J., Emmorey, K., Smith, J. F. & Braun, A. R. Symbolic gestures and spoken language are processed by a common neural system. Proceedings of the National Academy of Sciences 106, 20664–20669 (2009).

32. Allen, E. J. et al. A massive 7T fMRI dataset to bridge cognitive neuroscience and artificial intelligence. Nat Neurosci 25, 116–126 (2022).

33. Stringer, C. et al. Spontaneous behaviors drive multidimensional, brainwide activity. Science 364, eaav7893 (2019).

34. Arcaro, M. J., Schade, P. F. & Livingstone, M. S. Body map proto-organization in newborn macaques. Proc Natl Acad Sci U S A 116, 24861–24871 (2019).

35. Arcaro, M. J. & Livingstone, M. S. On the relationship between maps and domains in inferotemporal cortex. Nat Rev Neurosci 22, 573–583 (2021).

36. Botvinick, M. & Cohen, J. Rubber hands ‘feel’ touch that eyes see. Nature 391, 756–756 (1998).

37. Yamamoto, S. & Kitazawa, S. Reversal of subjective temporal order due to arm crossing. Nat Neurosci 4, 759–765 (2001).

38. Roder, B., Kusmierek, A., Spence, C. & Schicke, T. Developmental vision determines the reference frame for the multisensory control of action. Proceedings of the National Academy of Sciences 104, 4753–4758 (2007).

39. Aqil, M., Knapen, T. & Dumoulin, S. O. Divisive normalization unifies disparate response signatures throughout the human visual hierarchy. Proc Natl Acad Sci U S A 118, e2108713118 (2021).

40. Kay, K. N., Winawer, J., Mezer, A. & Wandell, B. A. Compressive spatial summation in human visual cortex. Journal of Neurophysiology 110, 481–494 (2013).

41. Haggard, P., Christakou, A. & Serino, A. Viewing the body modulates tactile receptive fields. Exp Brain Res 180, 187–193 (2007).

42. Reed, C. L. & Farah, M. J. The psychological reality of the body schema: A test with normal participants. J Exp Psychol Hum Percept Perform 21, 334–343 (1995).

43. Wold, A., Limanowski, J., Walter, H. & Blankenburg, F. Proprioceptive drift in the rubber hand illusion is intensified following 1 Hz TMS of the left EBA. Front Hum Neurosci 8, 390 (2014).

44. Pulvermuller, F., Hauk, O., Nikulin, V. V. & Ilmoniemi, R. J. Functional links between motor and language systems. Eur J Neurosci 21, 793–797 (2005).

45. Finn, E. S. & Bandettini, P. A. Movie-watching outperforms rest for functional connectivity-based prediction of behavior. NeuroImage 235, 117963 (2021).

46. Sonkusare, S., Breakspear, M. & Guo, C. Naturalistic Stimuli in Neuroscience: Critically Acclaimed. Trends in Cognitive Sciences 23, 699–714 (2019).

47. Robertson, C. E. & Baron-Cohen, S. Sensory perception in autism. Nat Rev Neurosci 18, 671–684 (2017).

48. He, J. L. et al. A working taxonomy for describing the sensory differences of autism. Molecular Autism 14, 15 (2023).

49. Volkmar, F. R. & Nelson, D. S. Seizure Disorders in Autism. Journal of the American Academy of Child & Adolescent Psychiatry 29, 127–129 (1990).

50. Lee Masson, H., et al. Intact neural representations of affective meaning of touch but lack of embodied resonance in autism: A multi-voxel pattern analysis study. Molecular Autism 10, 39 (2019).

## References for Methods

1. Ugurbil, K., et al. Pushing spatial and temporal resolution for functional and diffusion MRI in the Human Connectome Project. Neuroimage 80, 80–104 (2013).

2. Glasser, M. F. et al. The minimal preprocessing pipelines for the Human Connectome Project. Neuroimage 80, 105–124 (2013).

3. Gershon, R. C. et al. Language Measures of the NIH Toolbox Cognition Battery. J Int Neuropsychol Soc 20, 642–651 (2014).

4. 1200 Subjects Data Release - Connectome. https://www.humanconnectome.org/study/hcp-young-adult/document/1200-subjects-data-release.

5. Robinson, E. C. et al. MSM: A new flexible framework for Multimodal Surface Matching73. Neuroimage 100, 414–426 (2014).

6. Glasser, M. F. et al. A multi-modal parcellation of human cerebral cortex. Nature 536, 171–178 (2016).

7. Glasser, M. F. & Van Essen, D. C. Mapping human cortical areas in vivo based on myelin content as revealed by T1- and T2-weighted MRI. J Neurosci 31, 11597–11616 (2011).

8. Haak, K. V. et al. Connective field modeling. Neuroimage 0, 376–384 (2013).

9. Benson, N. C. & Winawer, J. Bayesian analysis of retinotopic maps. eLife 7, e40224 (2018).

10. Saadon-Grosman, N., Loewenstein, Y. & Arzy, S. The ‘creatures’ of the human cortical somatosensory system. Brain Commun 2, fcaa003 (2020).

11. Ma, S. et al. An fMRI dataset for whole-body somatotopic mapping in humans. Sci Data 9, 515 (2022).

12. Reuter, M., Wolter, F.-E. & Peinecke, N. Laplace–Beltrami spectra as ‘Shape-DNA’ of surfaces and solids. Computer-Aided Design 38, 342–366 (2006).

13. Reuter, M., Biasotti, S., Giorgi, D., Patane, G. & Spagnuolo, M. Discrete Laplace–Beltrami operators for shape analysis and segmentation. Computers & Graphics 33, 381–390 (2009).

14. la Tour, T. D., Eickenberg, M., Nunez-Elizalde, A. O. & Gallant, J. L. Feature-space selection with banded ridge regression. Neuroimage 264, 119728 (2022).

15. Hoerl, A. E. & Kennard, R. W. Ridge Regression: Biased Estimation for Nonorthogonal Problems. Technometrics 12, 55–67 (1970).

16. Wang, L., Mruczek, R. E., Arcaro, M. J. & Kastner, S. Probabilistic Maps of Visual Topography in Human Cortex. Cereb Cortex 25, 3911–3931 (2015).

17. Allen, E. J. et al. A massive 7T fMRI dataset to bridge cognitive neuroscience and artificial intelligence. Nat Neurosci 25, 116–126 (2022).

18. Singmann, H. et al. Afex: Analysis of Factorial Experiments (2024).

19. Lenth, R. V., et al. Emmeans: Estimated Marginal Means, aka Least-Squares Means (2024).

20. Burt, J. B., Helmer, M., Shinn, M., Anticevic, A. & Murray, J. D. Generative modeling of brain maps with spatial autocorrelation. NeuroImage 220, 117038 (2020).

21. Lin, T.-Y. et al. Microsoft COCO: Common Objects in Context. arXiv:1405.0312 [cs] (2015).

22. Washington-University/workbench: Connectome Workbench. https://github.com/Washington-University/workbench.

23. Cao, Z., Hidalgo, G., Simon, T., Wei, S.-E. & Sheikh, Y. OpenPose: Realtime Multi-Person 2D Pose Estimation using Part Affinity Fields (2019). 1812.08008.

